# The kinetic landscape of human transcription factors

**DOI:** 10.1101/2022.06.01.494187

**Authors:** Nicholas E Mamrak, Nader Alerasool, Daniel Griffith, Alex S Holehouse, Mikko Taipale, Timothée Lionnet

## Abstract

Cell-to-cell variability is shaped by transcription dynamics because genes are transcribed in bursts interspersed with inactive periods. The stochasticity of bursting means that genes transcribed in rare bursts exhibit more heterogeneity at the single cell level than genes that burst often ^1, 2^. Transcription starts with the binding of Transcription Factors (TFs) to specific sequence motifs where they recruit the transcription machinery ^3^. In some systems, individual TF binding events temporally correlate with the firing of transcriptional bursts, defining the target gene’s frequency and duration ^4–6^. However, in the absence of methods that assess the impact of different TFs on transcription dynamics at the same genetic loci, it remains unclear whether DNA binding kinetics are the sole determinant of bursting. Here we develop an imaging-based synthetic recruitment assay, CRISPRburst, and measure how 92 human TFs impact bursting kinetics. We show that TFs recruited to chromatin under identical conditions generate diverse bursting signatures, some TFs increasing the probability of the gene turning on while others increase the number of mRNA molecules transcribed per burst. We find that the association of TFs with specific protein partners determines their bursting output, and train a model to predict the kinetic signatures of all human TFs. These kinetic signatures can be used as a TF classification system complementary to existing families based on DNA binding domains. Additionally, kinetic signatures provide a rational framework to design synthetic activators, model transcription regulation, and understand expression heterogeneity.

## Introduction

Transcription factors (TFs) regulate changes in gene expression in response to ever-changing cellular environments. Genome-wide assays mapping TF occupancy across the genome have uncovered a wide range of sequence motifs ^7^ and abilities to access closed chromatin ^8, 9^. While chromatin occupancy is a clear regulator of TF function, it remains unclear how TFs vary in their ability to induce transcription once bound.

Live imaging and single-cell techniques demonstrate that transcription is inherently discontinuous ^10–12:^ individual alleles exhibit bursts of activity interspersed by inactive periods. The simplest description of bursting kinetics requires three parameters: the frequency at which bursts are initiated, their duration, and intensity (how many RNA molecules are transcribed per burst) ^13^. Small frequent bursts or large rare bursts can give rise to identical expression levels on average, but these regulatory strategies promote low and high heterogeneity across a cell population, respectively. While cell-to-cell variability is increasingly appreciated as a regulator of biological processes ranging from development and aging to cancer and viral infection ^14–18^, the molecular drivers of bursting and expression heterogeneity remain incompletely understood.

It has been proposed that bursts are directly linked to TF binding events ^4–6, 10^. In this model, the on- and off-rates of a TF to regulatory sequences define the frequency and (inverse) duration of bursts, while the strength of the TF activation domain governs the mRNA firing rate during a burst. While this model is appealing, some systems exhibit complex dynamics, which suggest extra regulatory layers^13, 19–21^. For instance, pluripotency TFs remain bound to enhancers long after their target genes stop expressing during early differentiation ^22^. In the absence of techniques able to decouple TFs activation strength from their occupancy, which kinetic step(s) are regulated by TFs remains unclear.

Here we present CRISPRburst, a CRISPR/Cas9-mediated TF recruitment and fluorescence imaging framework, and use it to characterize how 92 TFs impact transcription kinetics. Overall, TFs can modulate both the probability of a gene being active and the intensity during these active periods. While most individual TFs act only on one of these two kinetic metrics, a small number of ‘multi-tasking’ TFs affect both. TFs with similar kinetic signatures share interactions with co-activating complexes, but differ in features typically used to classify TFs, such as DNA binding domain homology. We use the CRISPRburst dataset to train a multivariate regression model able to predict the functional effects of 2,300 human TFs and co-activators, providing a systems-level view of the functional diversity of human transcription factors.

## Results

### CRISPRburst, an inducible dCas9-mediated recruitment platform to study transcription kinetics

In order to assay the activating effects of TFs while bypassing their differences in DNA binding affinity, we turned to a synthetic recruitment strategy (Figure 1A): we generated a cell line stably expressing nuclease-defective Cas9 (dCas9) fused to the dimerization domain ABI1. TFs fused to ABI1’s cognate domain PYL1 are recruited to the locus bound by ABI1-dCas9 upon the addition of phytohormone *S*-(+) - abscisic acid (ABA) ^23–25^, enabling precise spatiotemporal control. This strategy provides a straightforward control for non-specific effects of TF overexpression (comparing conditions with and without ABA), and scales easily because PYL1 is small compared to dCas9 (176 vs. 1360 amino acids ^23^), facilitating library generation.

**Figure 1:**
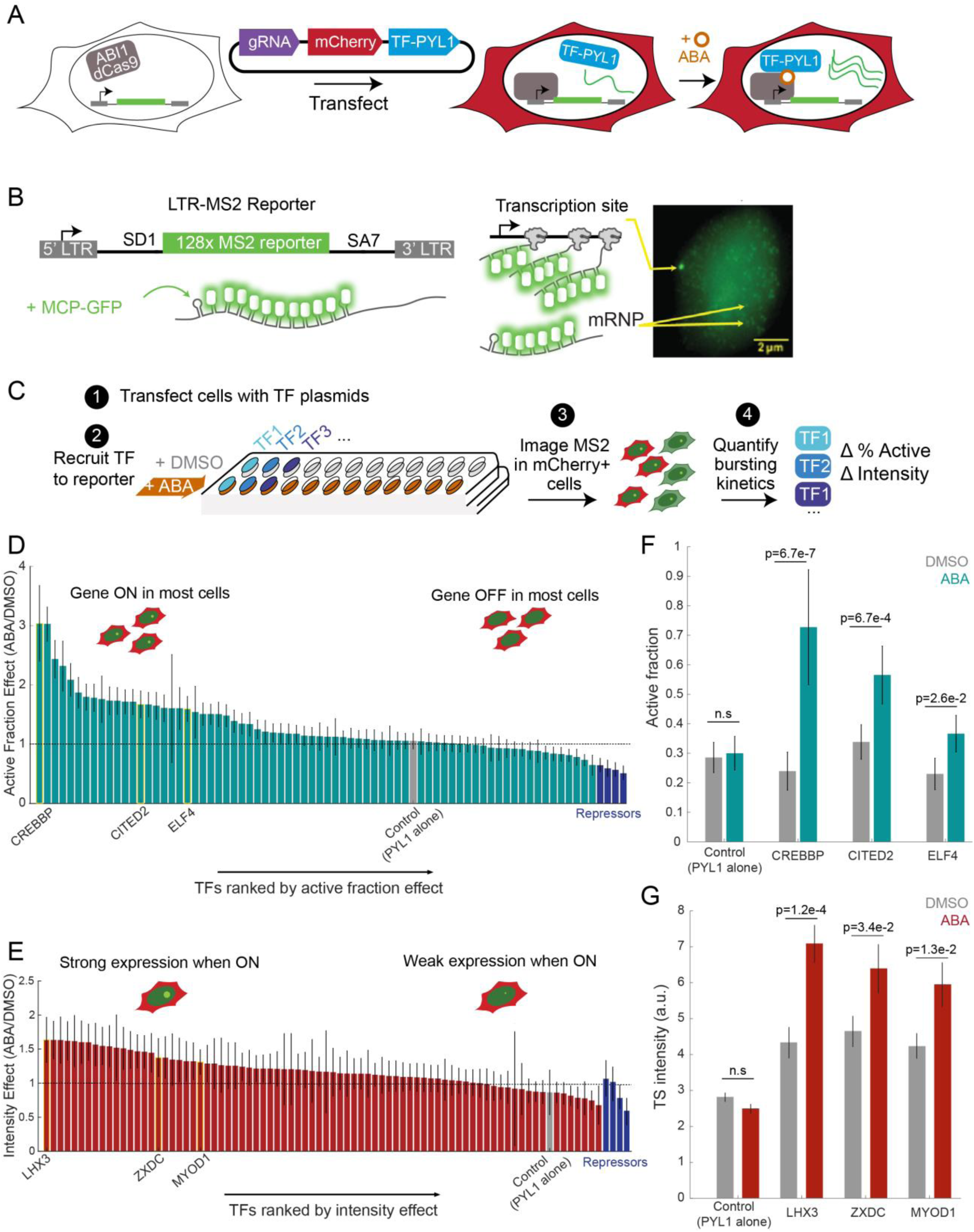
CRISPRburst enables comparing the impact of diverse TFs on transcription kinetics. **A** CRISPRburst principle. Cells stably expressing ABI1-dCas9 and a fluorescent mRNA reporter are transfected with a plasmid encoding a TF-PYL1 fusion, an sgRNA targeting the reporter promoter, and an mCherry transfection marker. Addition of abscisic acid (ABA) enables monitoring changes in transcription dynamics of the reporter gene upon recruitment of the TF-PYL1 fusion to promoter-bound dCas9. **B** Left: LTR-MS2 reporter gene. The MS2 cassette is inserted between splice donor (SD1) and splice acceptor (SA7) sites. Right: Example cell showing the MS2 reporter transcription site (bright nuclear spot). Several unspliced mRNA Particles (mRNPs) are visible in the nucleus, consistent with previous demonstration that splicing occurs post-transcriptionally in this system^29^. **C** Experimental timeline. LTR-MS2 cells stably expressing ABI1-dCas9 are seeded in 96 well plates and 1) transfected with plasmid encoding a TF-PYL1 fusion, gRNA targeting the LTR, and an mCherry marker. 2) TFs are recruited to the LTR-MS2 reporter using ABA. 3) Live cells are imaged 16 h post-recruitment. 4) mCherry positive cells are analyzed for quantification of transcription site activity and intensity. **D, E** Transcription factors ranked by decreasing active fraction effect (the fraction of cells with an active reporter transcription site upon TF recruitment (ABA) over the fraction when the TF is not recruited (DMSO), **D**) and decreasing intensity effect (mean transcription site intensity upon TF recruitment (ABA) over mean intensity when the TF is not recruited (DMSO), **E**). PYL1 Control: gray; KRAB repressors: dark blue. **F, G** Example TFs demonstrate the range of active fraction (**F**) and intensity effects (**G)** in the absence of (gray) or upon TF recruitment (teal or red). An average of 220 cells were analyzed per TF. Significance was determined by two sample t-tests. Error bars mark the standard error on the mean.

The HIV 5’ long terminal repeat (LTR) is an important model because stochastic HIV transcription underlies cycles of activity and latency that prevent full eradication of the virus from infected patients ^18, 26–28^. In order to quantify HIV transcription dynamics, we leveraged an established cell line containing a single integration of the human immunodeficiency virus 1 (HIV-1) genome in which the viral open reading frames are replaced with an intronic 128x MBS (MS2 Binding Site) cassette for fluorescent mRNA labeling ^29^ (Figure 1B). When transcribed, MBS sequences become bound by MCP (MS2 Capsid Protein) fused to GFP, enabling reliable detection of nascent mRNA in real time in living cells ^29, 30^.

In total, the LTR-MS2 cell line stably expresses 1) the LTR-MBS reporter gene, 2) the MCP-GFP protein that binds nascent mRNAs, and 3) ABI1-dCas9 (Figure 1A). In order to mimic the latent state, the cell line does not express the HIV trans-activator Tat, which could otherwise confound results via interactions with Positive Transcription Elongation Factor b (P-TEFb) ^31^. TFs are then assayed by transfecting LTR-MS2 cells with an arrayed library of plasmids encoding both a TF-PYL1 fusion and the single guide RNA (sgRNA) targeting the HIV LTR. The sgRNA target sequence overlaps with a naturally occurring NF-κB binding site in the HIV LTR ^32^ (Figure S1A), ensuring that recruited TFs occupy a physiologically relevant position relative to nucleosomes and the transcription start site ^33^.

The high signal-to-noise ratio afforded by the bright MS2 reporter enables nascent transcription detection in multi-well plates using a 20x magnification high content screening platform (Note 1, Figure 1C). In each condition, we quantified both 1) the active fraction, i.e. the probability that the reporter is active in each cell, and 2) the fluorescence intensity of the transcribing spot (directly proportional to the number of nascent transcripts at the reporter ^29^). We then normalized metrics computed when each TF is recruited (ABA) to their corresponding non-recruited (DMSO) conditions in order to isolate effects specific to TF recruitment.

### Functional characterization of TFs using an imaging-based synthetic recruitment assay

Applying CRISPRburst, we characterized 78 human TFs known to induce transcription through CRISPR activation (CRISPRa) ^25^. Those include sequence-specific TFs spanning several families, transcriptional co-activators, and a synthetic activator (VP64). We also included 4 KRAB (Krüppel associated box) containing repressors, and the PYL1 moiety alone as controls.

Upon recruitment, 28 TFs generate an increase in reporter active fraction (ABA/DMSO ratio > 1.30; Figure 1D) and 40 TFs generate an increase in reporter intensity (ABA/DMSO ratio > 1.15; Figure 1E). Thus, both the fraction of the time the gene is active and the intensity of active periods can be tuned by TFs. As expected, known strong activators such as the histone acetyltransferases CREBBP and p300, and the synthetic activator VP64 exhibited large increase in the active fraction, (Figure 1D, Table S1), while KRAB repressors exhibited the lowest active fraction effect ^34^. The TFs exhibiting the highest intensity effects were LIM/Homeobox protein LHX3, globin transcription factor GATA1, and cyclin-dependent kinase CDK9 (Figure 1E, Table S1). Overall, TF effect sizes range from 0.64 to 3.04 for active fraction and 0.68 to 1.64 for intensity (Figure 1F-G, S1E), suggesting that the frequency and/or duration of active periods provides a wider regulatory range than burst intensity.

The baseline active fraction of our reporter is 0.3 (Figure 1F), meaning an effect size of 3 for active fraction is approaching the theoretical maximum (active fraction = 1.0). The maximal intensity per transcription site (TS) is likely limited by physical constraints of the transcription machinery as a limited number of RNA polymerase molecules can be loaded per gene due to polymerase velocity and spacing ^29^. The ranges of effect sizes observed here in nascent transcription levels lead to dramatic changes in protein expression, likely due to downstream amplification steps (Figure 5C). We controlled that differences between TFs do not reflect variable transfection efficiency (Figure S1F-G).

If the frequency and duration of active periods were solely defined by TF binding, one would expect that TFs recruited via dCas9 would all exhibit similar active fractions and vary only in their burst intensity. The diverse range of active fractions and burst intensities induced by dCas9-recruited TFs therefore clearly indicates a more complex regulatory architecture.

To ensure that TFs exhibit similar behaviors when recruited by dCas9 as they do in physiological conditions, we verified that 6 factors known to directly bind the HIV-1 5’ LTR ^35^ regulate similar kinetic steps upon overexpression alone (DMSO) as they do when recruited by dCas9 (ABA). Indeed, the effects observed in both conditions (Figure S1H-I) are positively correlated (Pearson’s Correlation Coefficient, PCC = 0.50 and 0.84 for active fraction and intensity, respectively), and more so than those of TFs that do not bind the LTR (PCC = 0.42 and 0.72).

### Bursting kinetics define distinct TF classes

We observed that TFs preferentially impact either the probability of the reporter being active, or the intensity of gene when on (Figure 2A-C), suggesting that individual TFs have evolved specialized kinetic roles. Consistent with this idea, the majority of TFs sampled fall in two main classes upon clustering (see methods, Figure 2D, Figure S2A-D, Table S2): 1) TFs increasing the active fraction alone, and 2) TFs increasing the intensity alone. The rest of the TFs fall in two smaller classes containing ‘multi-tasking’ TFs that impact both the active fraction and intensity, and a final class of TFs with little impact on either parameter.

**Figure 2:**
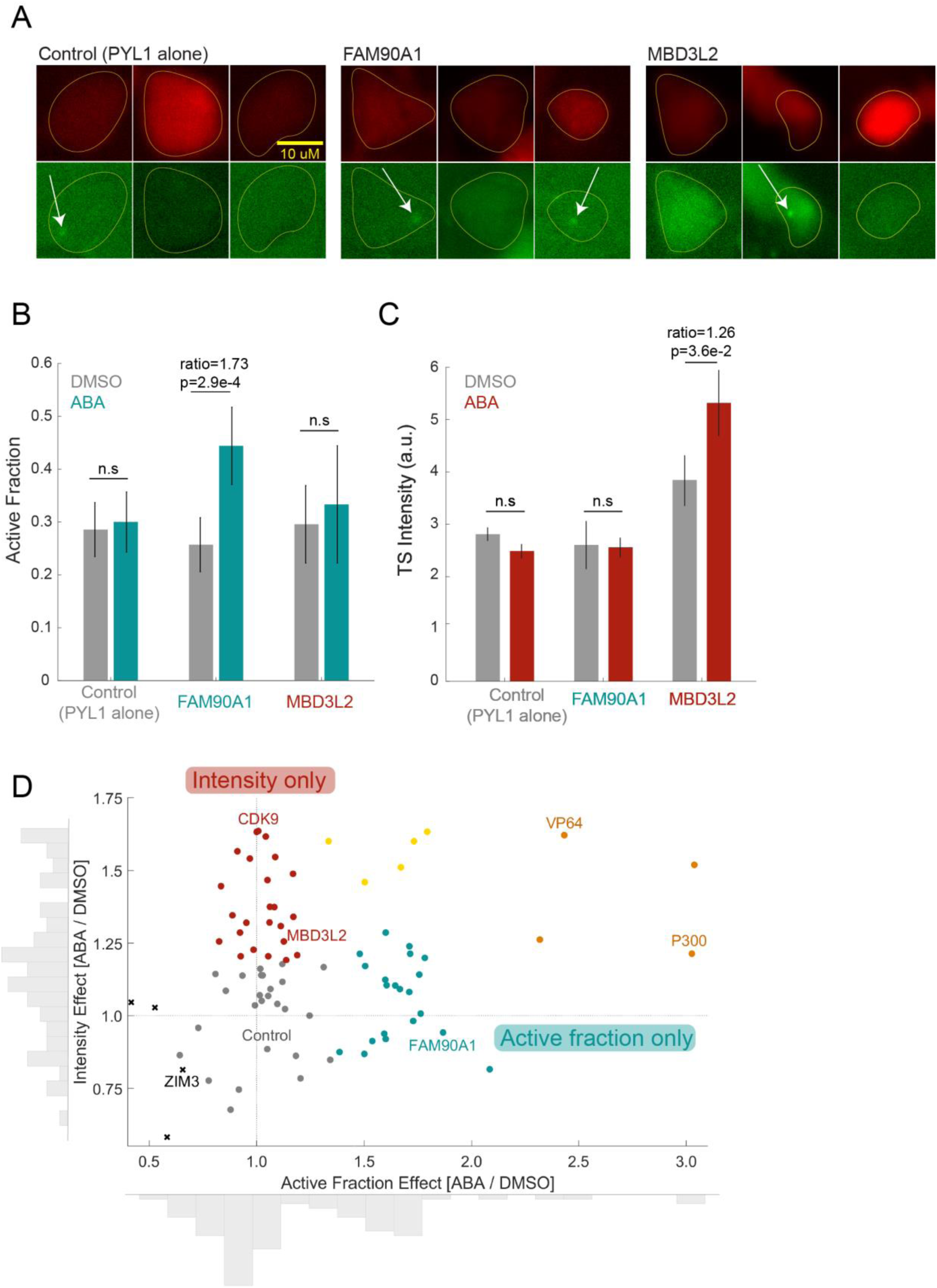
TFs have diverse effects on transcription kinetics. **A** Representative images of PYL1, FAM90A1, and MBD3L2 transfected cells upon TF recruitment. Top: mCherry (transfection marker). Bottom: MCP-GFP. Yellow outlines: mCherry mask. White arrows: transcription sites. Scale bar: 10 μm. **B, C** FAM90A1 selectively increases the active fraction (**B**) but not the intensity (**C**) of the reporter, while MBD3L2 exhibits the opposite specialization. Non-recruited TF (DMSO): gray, recruited TF (ABA): teal or red. **D** Active fraction versus intensity for 78 characterized TFs. Colors mark the cluster each TF belongs to (see text). Teal represents active fraction only TFs, red represents intensity only TFs, yellow and orange represent low and high strength multi-taskers, and gray represents TFs with low effect. Black Xs: KRAB repressors. Significance was determined by two sample t-tests. Error bars designate standard error on the mean.

The activation domains of TFs often lie within intrinsically disordered regions (IDRs) that can mediate the formation of hubs (dynamic non-stoichiometric molecular assemblies associated with sites of active transcription, also referred to as transcriptional condensates) ^36^. Since hub formation and other biophysical properties are tuned by the length and amino acid content of IDRs ^37, 38^, we hypothesized that TFs from different kinetic clusters might contain IDRs with distinct sequence biases. Indeed, TFs in multi-tasking clusters contain cumulatively longer IDRs compared to TFs with little effect (p = 2.5 x 10^-2^, Figure S3A), and TFs increasing either kinetic metric are enriched for certain amino acids consistent with known biases of IDRs ^37, 39–41^ (Table S3, Figure S3B). However, we did not detect significant differences between IDR features of TFs affecting only the active fraction and those of TFs affecting only the intensity (Figure S3C-D). Together, these findings suggest that while the biophysical properties of IDRs may tune the amplitude of TFs’ effects, they likely do not solely encode TF kinetic specialization.

### Interactions with co-activators are more predictive of TF kinetic specificity than IDR features

As most TFs do not have intrinsic catalytic activity and rather regulate transcription by recruiting co-activators, we reasoned that kinetic specialization could stem from the preferential recruitment of distinct co-activator subsets (Figure 3A, top). To test this hypothesis, we built a model predicting TF kinetic signatures based on the co-activators they bind. We collected interaction partners for each human TF from the Biological General Repository for Interaction Datasets (BioGRID) ^42^ and assigned a predictive weight to each possible interactor in a partial least-squares multivariate regression model. This model uses these weights to predict an active fraction and intensity effect for over 2,300 human TFs (including co-activators, Figure 3A, bottom; Note 2; Table S5). Predictions increase in accuracy with increasing training set size (Figure S4), and show little sensitivity to BioGRID coverage (Figure S5A,C). We predicted the kinetic effects of each TF in our dataset based on training the model on all the other characterized TFs, and verified that the model accurately determines whether each TF increases the active fraction or intensity (Figure 3B, left).

**Figure 3:**
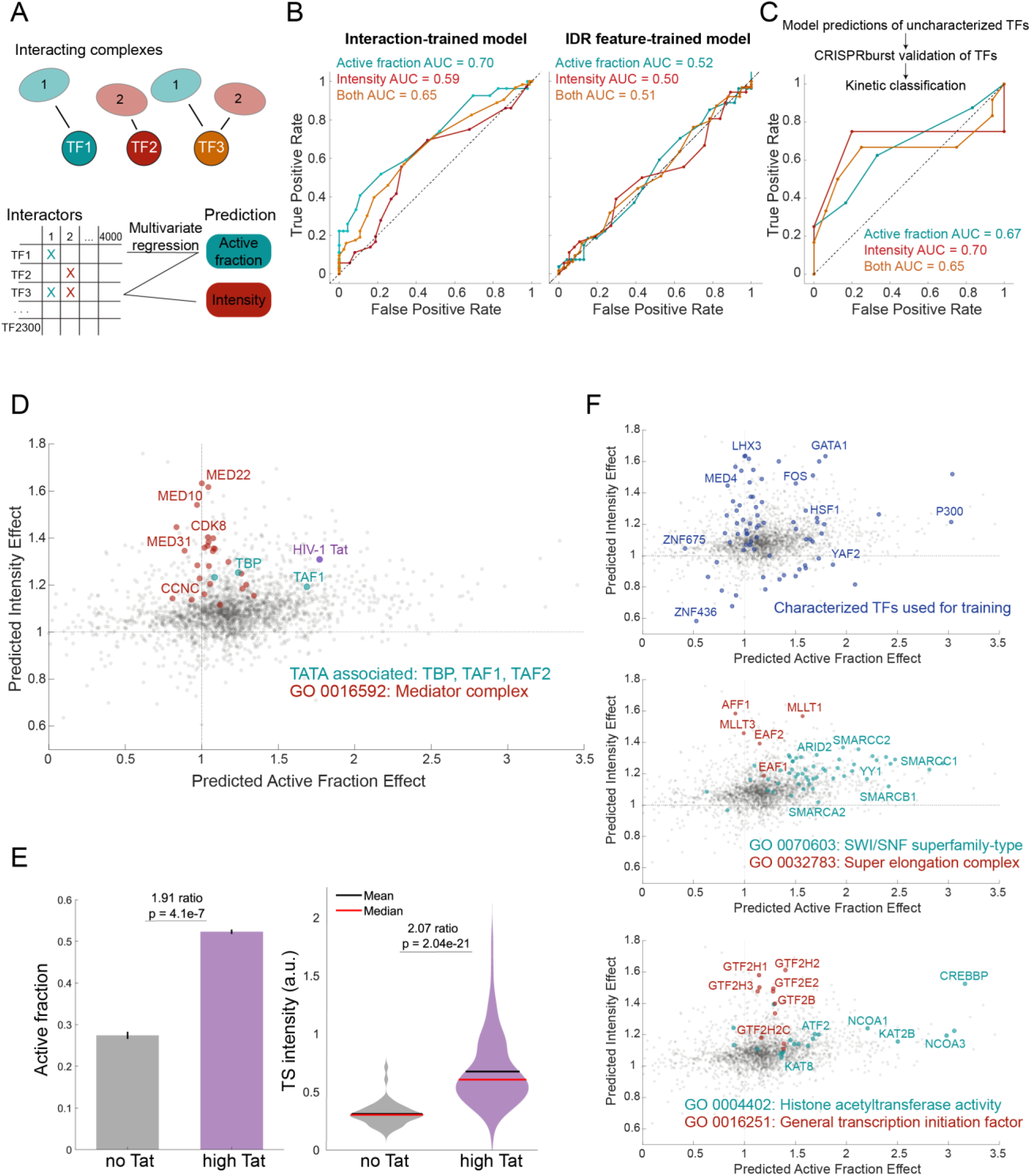
An interaction-based model predicts the kinetic landscape of human TFs. **A** Top: TF association with distinct co-activators, biases their kinetic role. Bottom: the model links TF interactions to bursting kinetics through multivariate regression. **B** Receiver operating characteristic (ROC) evaluation of the model’s ability to classify TFs as increasing the active fraction or intensity. We predicted the active fraction and intensity of each TF in our training set using a model trained on all other TFs. Model trained on interaction data (left) or IDR features (right). **C** ROC evaluation as in **B** applied to 14 TFs never seen by the model. **D** Predicted kinetics of HIV-1 trans-activating protein Tat (purple) overlaid with predictions for all human TFs (gray). Teal: TATA associated factors (TBP, TAF1, TAF2); red: GO term 0016592 for Mediator complex associated factors. **E** Active fraction (left) and Intensity (right) measurements in a cell line not expressing Tat (gray) vs. a cell line expressing high Tat levels. An average of 200 cells were analyzed per condition. Significance was determined by two sample t-tests. **F** The kinetic landscape of human transcription factors highlighting Top: the TFs used for model training data (dark blue); Middle: GO term 0070603 SWI/SNF superfamily-type associated factors (teal) and GO term 0032783 Super elongation complex factors (red); Bottom: GO term 0004402 histone acetyltransferase activity associated factors (teal) and GO term 0016251 RNAPII general transcription initiation factors (red). Error bars designate standard error on the mean.

We went on to profile by CRISPRburst 14 TFs never seen by the model, and verified that the model assigns the TFs from the validation set into the correct kinetic class with 71.4% accuracy and 75.0% specificity (Figure 3C, S5E-G, Table S4). In parallel, we built a model predicting TF signatures using IDR features as training data in which weights are assigned to each amino acid frequency as well as length, complexity, hydrophobicity, aromaticity, and charge distribution. This model was unable to classify TFs into kinetic classes (Figure 3B, right), demonstrating that TF-cofactor interactions play a greater role in specifying kinetic function than IDR sequence content.

Does the CRISPRburst-based model predict the behavior of TFs upon physiological recruitment? The model predicts that Mediator subunits should increase the reporter intensity while TATA-binding protein (TBP) should increase both active fraction and intensity (Figure 3D). Indeed, studies using the same HIV-1 LTR model show that Mediator increases Pol II re-initiation, leading to more intense bursts, while TBP regulates the timing of active and inactive periods and the number of nascent transcripts per transcription site through initiation rate ^29, 43^. To further confirm that predictions apply to TFs in their normal context, we turned to the viral trans-activator, Tat, predicted by the model to increase both active fraction and intensity (Figure 3D). We compared the active fraction and intensity of the reporter between cell lines expressing either high levels of Tat or no Tat. Consistent with model predictions, high Tat cells exhibited a significant increase in both the reporter active fraction (p = 4.1 x 10^-7^) and intensity (p = 2.0 x 10^-21^) relative to their No Tat counterparts (Figure 3E). Together, these data validate that TF kinetic roles generalize beyond the context of the synthetic recruitment assay.

### Kinetic signatures map to regulatory steps and specialized co-activators

Our model predictions provide a kinetic landscape of human transcription factors and co-activators (Figure 3F). Specific complexes or functional families cluster in distinct regions: Mediator subunits are biased for strong intensity, consistent with Mediator’s role in rapid re-initiation ^29, 44^, while histone acetyltransferases exhibit strong active fraction, in line with their role in creating a chromatin environment permissive for transcription ^30, 45, 46^. Similarly, Super Elongation Complex associated TFs and general transcription factors increase burst intensity, while SWI/SNF subunits increase active fraction, both consistent with their known functions regulating RNA polymerase pausing and chromatin remodeling, respectively, but, have not been ascribed to specific kinetic features (Figure 3F). These observations are further supported by enrichment analysis (Figure 4A-B, Table S6).

**Figure 4:**
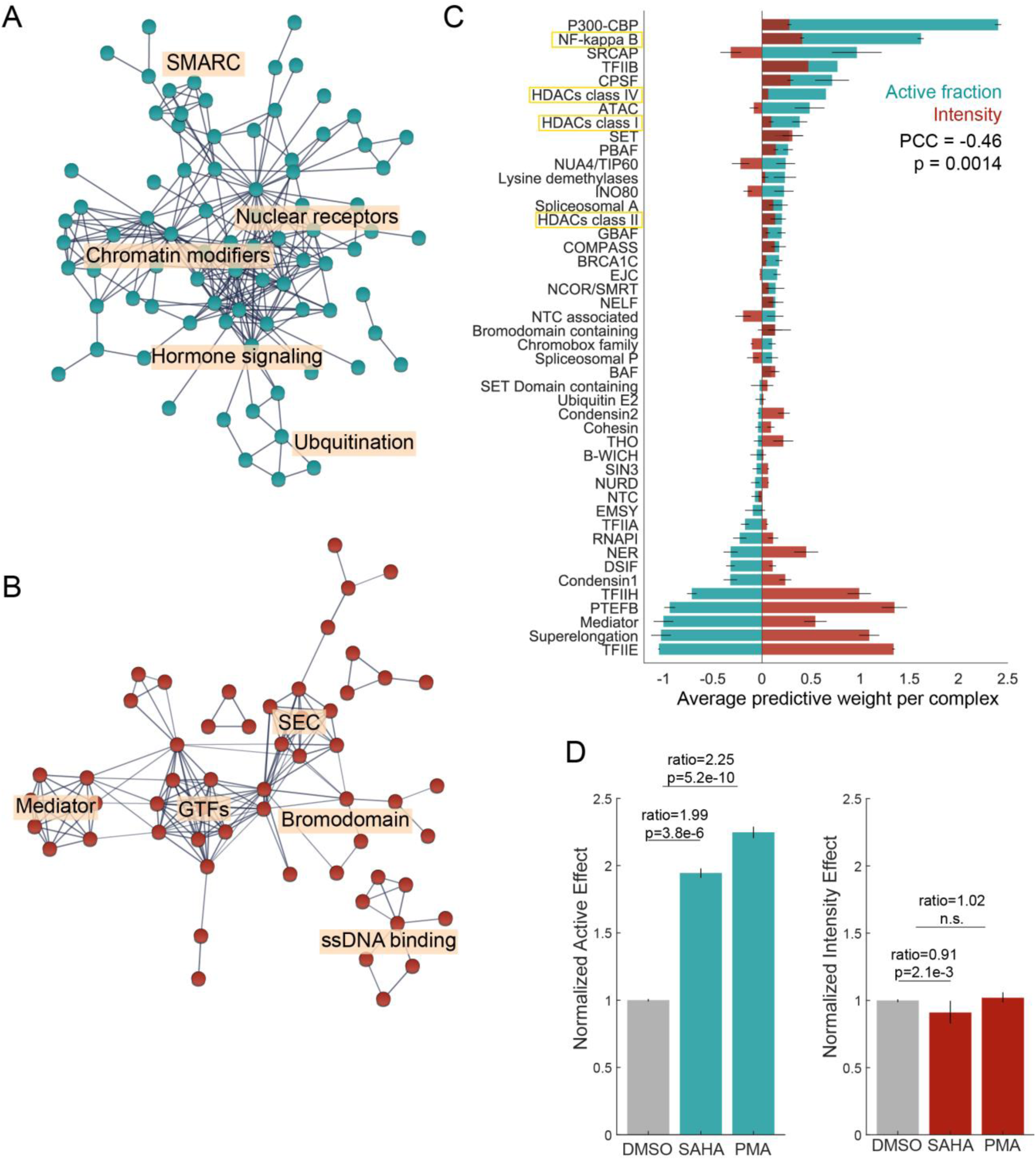
Co-activating complexes drive TF kinetic specialization. **A**, **B** Protein-protein interaction networks among the 100 interactors with highest predictive weights for active fraction (**A**) and intensity (**B**), created using STRING-db ^66^. Disconnected nodes are hidden for clarity. **C** Average weights per complex in the model predicting active fraction (teal) and intensity (red). PCC, Pearson’s correlation coefficient. **D** Small molecule inhibition of candidate active fraction regulators in LTR-MS2 reporter cells. Cells were treated with the HDAC inhibitor Vorinostat (SAHA), or small molecule activator of NF-κB, phorbol myristate acetate (PMA). Left, active fraction (normalized to DMSO). Right, intensity (normalized to DMSO). A minimum of 2,700 cells were analyzed per condition. Significance was determined by two sample t-tests. Error bars designate standard error on the mean.

To better understand the contributions of co-activators, we computed the mean predictive weight of known complexes (Figure 4C). Weights for active fraction and intensity effects are negatively correlated (PCC = -0.46, p = 0.0014), suggesting that co-activating complexes preferentially impact a single kinetic feature. While the weights likely translate the direct effects of co-activator recruitment, they may also reflect statistical associations, explaining why some co-activators are weighted negatively (e.g. it might be unusual for TFs binding P-TEFb to also bind p300/CREBBP, therefore these TFs end up less likely to increase the active fraction), or why complexes associated with repressive functions exhibit positive predictive weights (TFs increasing the active fraction might be more likely to be repressed via HDAC association).

To validate the kinetic specializations assigned to specific complexes, we treated the LTR-MS2 reporter cells with the histone deacetylase (HDAC) inhibitor Vorinostat (SAHA), or a small molecule activator of NF-κB, phorbol myristate acetate (PMA). Both HDACs and NF-κB exhibit high weights for active fraction (Figure 4C). Indeed, treatment with 500 nM SAHA or 1uM PMA for 48 h increased the fraction of active reporter genes (ratios = 1.99 and 2.25, p = 3.8 x 10^-6^ and 5.2 x 10^-10^ for SAHA and PMA, respectively; Figure 4D) without substantially changing their intensity. Taken together, these results support that differential interactions of TFs with specialized co-activators shape the TF kinetic landscape.

### TF families exhibit broad kinetic diversity

TFs are traditionally classified into families sharing homologous DNA binding domains ^7^. The model predicts that kinetic variability within each TF family is much greater than the variability between family-averaged kinetics (Figure S6A). One exception is the KRAB domain-containing family, overwhelmingly predicted to be repressors. Interestingly, the family-defining KRAB domain does not bind DNA but recruits cofactors, consistent with the idea that DNA binding domains provide little information on kinetic specialization (Figure S6B).

### Oncogenes are biased for TFs increasing the active fraction

TF dysregulation drives many cancers. While oncogenes are enriched in regulators of specific pathways such as DNA repair or differentiation, it remains unclear whether the oncogenic potential of a TF is determined by features other than the function of its targets ^47^. Using the model, we predicted the effect on active fraction and intensity of transcription regulators commonly mutated in cancer, based on the catalog of somatic cancer mutations (COSMIC) ^48^. Mutated genes are enriched for TFs increasing the active fraction over those increasing intensity (p = 2.9 x 10^-11^; Figure S7A). This enrichment holds for TFs commonly found in oncogenic fusions^49^ (Figure S7B). Furthermore, the likelihood of TFs to act as driver genes correlates with the strength of their active fraction effect (p = 5.7 x 10^-3^; Figure S7C), but not their intensity^50^.

Lastly, we predicted the kinetic effects of TFs with expression levels associated with favorable or unfavorable survival across several cancer types ^51^. We found these prognostic TFs exhibited significant differences in active fraction strength, but not intensity, between favorable and unfavorable groups across a number of cancer types (Figure S7E). Some cancer types displayed associations between stronger active fraction effects and unfavorable outcomes, while others displayed an association with favorable outcomes. This is likely a reflection of the diversity of mechanisms in which tumors can dysregulate transcription. Collectively, these findings link the oncogenic potential of TFs with their ability to activate genes *de novo* rather than hypertranscribing lowly expressed targets.

### Kinetically distinct TFs elicit distinct re-activation outcomes in a T-cell model of HIV-1 latency

HIV-infected cells escape immune recognition by entering latency, a long-lived state of silenced viral expression. Latency reversal strategies have the potential to clear reservoirs of latent cells, but current reversal agents lack specificity ^52^. We hypothesized that TFs modulating the active fraction in CRISPRburst would likely increase escape from latency, while TFs increasing the intensity alone should have little effect. To test this idea, we chose one TF from each class - JAZF1 (active fraction), LHX3 (intensity), and p300 (multi-tasker) and recruited them to the HIV-1 LTR in a T cell model of latency, J-Lat 10.6 (HIV-R7/E^−^/GFP) ^53^ (Figure 5A-B). This cell line harbors a full-length HIV provirus containing a GFP open reading frame in place of the *Nef* gene. Using fluorescence-activated cell sorting (FACS), we quantified the percentage of cells with GFP signal indicative of re-activated viral transcription and escape from latency. In the absence of recruitment, none of the TFs had any effect; neither did recruitment of PYL1 (Figure S8). Upon recruitment, JAZF1 and p300 exhibited substantially larger increases in GFP+ cells than LHX3 (Figure 5C), demonstrating that TFs regulating the active fraction constitute more potent latency reversal agents than those regulating the intensity.

**Figure 5:**
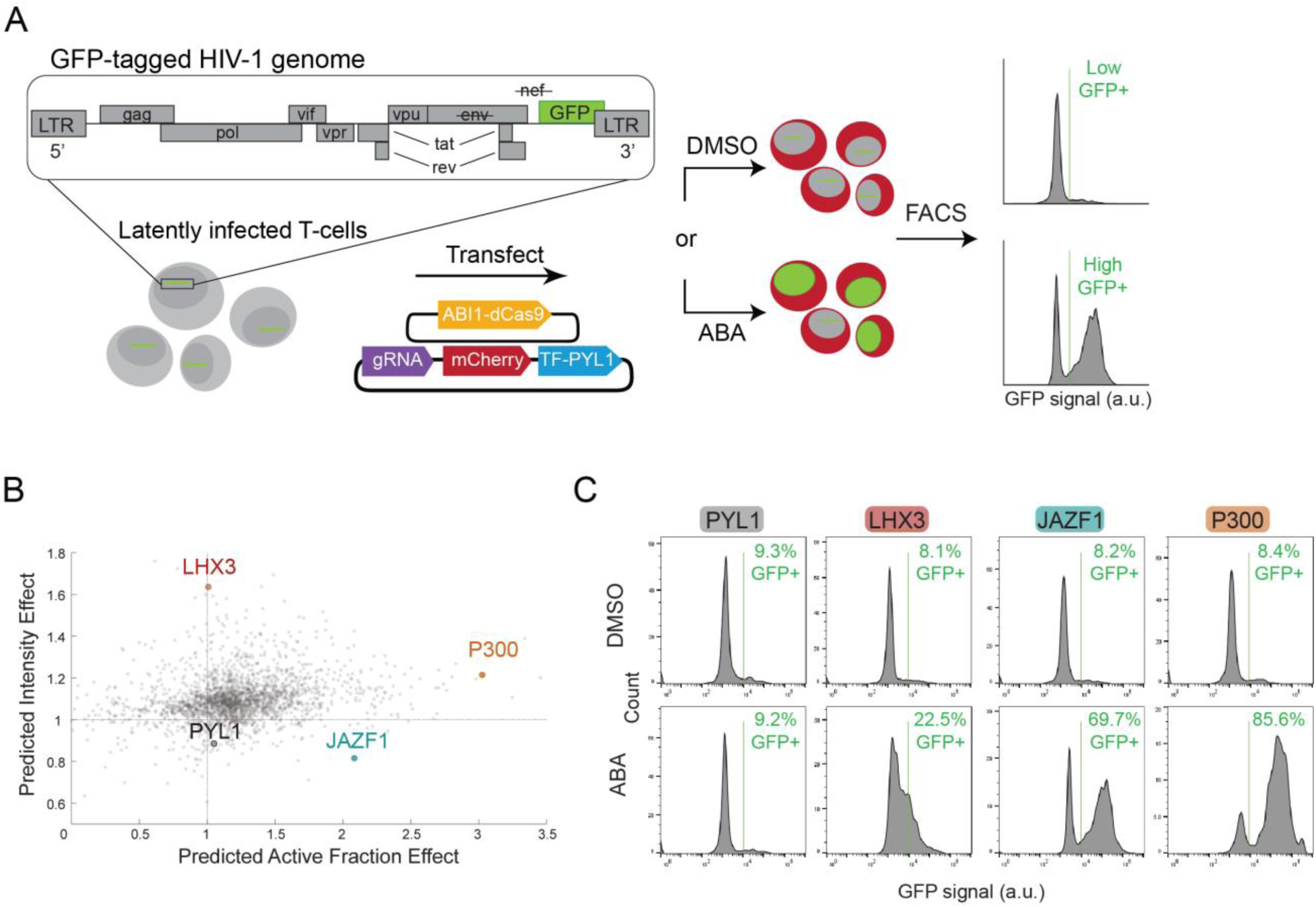
Specialized TFs elicit distinct re-activation outcomes in a T-cell model of HIV-1 latency. **A** Reactivation of HIV-1 provirus in a T-cell model of latency using specialized TFs. The reporter cassette (HIV-R7/E^−^/GFP) integrated into J-Lat 10.6 cells consists of a HIV provirus in which the *Nef* gene is replaced by GFP ^53^. J-Lat 10.6 cells were transfected with plasmids encoding ABI1-dCas9, sgRNA, TF-PYL1 fusion, and an mCherry marker. TFs are recruited to the LTR of HIV provirus using ABA. mCherry positive cells are analyzed via fluorescence-activated cell sorting (FACS) for GFP signal indicative of reactivation of viral transcription and escape from latency. **B** TF kinetic landscape, highlighting the three TFs used in J-Lat activation experiments. **C** FACS analysis of latency reactivation following TF recruitment to the HIV-1 LTR in J-Lat 10.6 cells. GFP+ frequencies of mCherry+ cells in the absence of (DMSO, top) or upon TF recruitment (ABA, bottom). A minimum of 215,000 cells were analyzed per sample.

## Discussion

DNA binding and occupancy have long been appreciated as a critical layer of transcription control ^29, 54, 55^; this study demonstrates that once bound, TFs control diverse kinetic outcomes based on co-activator choice. In line with the separation of tasks between DNA binding and activation originally proposed by Keegan and Ptashne ^56^, kinetic signatures provide a classification framework orthogonal and complementary to existing TF families based on DNA binding domains.

IDR-driven TF-TF association into dynamic hubs has recently emerged as a regulatory layer of the transcription process. Neither the active fraction nor the intensity correlates with specific amino acid bias or biophysical properties however, suggesting that TF kinetic specificities are unlikely to be solely determined by IDR sequence chemistry. However, TFs with longer IDRs are more likely to increase both the intensity and active fraction. This could suggest that long IDRs enable recruiting multiple co-activators; alternatively, they could potentiate the formation of hubs functioning as kinetic amplifiers.

While the architecture of the recruitment system, the nature of the promoter, or incomplete annotations of protein-protein interactions set a limit to the accuracy of kinetic predictions, we show that our model correctly assigns TFs into their kinetic families. Our study centered on the simple HIV promoter thus provides a robust conceptual framework to investigate more complex systems, e.g. how TFs synergize with one another ^7^, interact with core promoter motifs ^57^, or communicate to promoters from distal enhancers. This framework also provides direct avenues to rationally design next-generation synthetic activators, for instance building synergistic chimeras combining TFs from distinct kinetic classes.

## Methods

### Cell culture and treatments

LTR-MS2 reporter cells were a gift from Edouard Bertrand (IGMM, Fr). LTR-MS2 cells were maintained in Dulbecco’s Modified Eagle’s Medium (DMEM, Gibco 10569010) with 10% fetal bovine serum (FBS, VWR Seradigm 97068085), 1% penicillin-streptomycin (Gibco 15140122), and appropriate selection antibiotic when needed at 37°C with 5% CO2. Cells were routinely passed using Trypsin 0.25% EDTA dissociation reagent (Gibco 25200056). J-Lat 10.6 cells were a gift from Nathaniel Landau (NYU SoM). J-Lat cells were maintained in RPMI-1640 Medium (Gibco 11875085) with 10% FBS, 1% penicillin-streptomycin at 37°C with 5% CO2. For small molecule treatments, cells were treated with either 500 nM Vorinostat (SAHA, Sigma SML0061) or 1 μM phorbol myristate acetate (PMA, Sigma 79346) for 48 hours prior to imaging in LTR-MS2 imaging experiments or 1 μM PMA for 24 hours in FACS experiments using J-Lat cells.

### Plasmids

We created PYL1-TF fusion plasmids by LR recombination cloning using Gateway LR Clonase II enzyme mix (Invitrogen 11791020). The backbone PYL1 vector contains a CcdB toxin cassette that is replaced by the TF of interest upon successful recombination allowing for selection of colonies of interest ^25^. We transformed the recombination reaction products into CcdB-sensitive One Shot Stbl3 chemically competent cells (Invitrogen) and confirmed correct insertion by Sanger sequencing. We cloned the single-guide RNAs (sgRNAs) targeting the LTR-MS2 reporter into a modified pLCKO (Addgene #73311) to co-express guide RNAs from a hU6 promoter and mCherry from an hPGK promoter. We performed routine cloning using Q5 high-fidelity 2X master mix (NEB M0492) for PCR amplification and isothermal assembly using 2X Gibson master mix (NEB E2611) before transformation into NEB Stable or 5-alpha chemically competent cells.

### Virus production

We generated lentivirus by transiently transfecting 70% confluent HEK293T cells in 10cm plates with the plasmid encoding the construct of interest, as well as pMD2.G and psPAX2 ^58^ plasmids at a mass ratio of 2:1:1 (10 μg: 5 μg: 5 μg), using XtremeGENE HP (Roche) according to the manufacturer’s protocol. We collected virus-containing supernatant at 24 h, 48 h, and 72 h after transfection, pooled the supernatants and used one third of the resulting supernatant stock for each subsequent viral integration and creation of stably expressing cell lines.

### Stable cell line generation

We created ABI1-dCas9 stably expressing lines by infection with lentivirus particles generated from the pLX303 ABI1-dCas9 ^25^ plasmid which encodes a cassette co-expressing a blasticidin resistance gene from the hPGK promoter. We treated LTR-MS2 reporter cells placed in 10 cm dishes with viral supernatant supplemented with 8 μg/ml polybrene (Sigma TR1003G). 24 h later, we replaced the cell medium with fresh DMEM containing 8 μg/ml blasticidin to select LTR-MS2 reporter cells positive for ABI1-dCas9. After selection and expansion, we confirmed stable integration of ABI1-dCas9 using PCR genotyping.

### Reporter transcription site image acquisition

We plated 10,000 LTR-MS2 reporter cells stably expressing ABI1-dCas9 in each well of black, clear bottom 96 well plates (Thermo 165305). We transfected cells with 50 ng of PYL1-TF fusions and reporter-targeting sgRNA using XtremeGENE HP (Roche) according to the manufacturer’s protocol. Experiments were performed using either one plasmid co-expressing TF-PYL1 and mCherry or by co-transfecting separate plasmids, one expressing TF-PYL1 and one expressing mCherry. There was no significant relationship between mCherry expression and the observed TF effects using either method (Figure S1F-G). 24 hours post transfection, we treated cells with either 100 μM abscisic acid (ABA) or matched DMSO vehicle control for 16 h before imaging, in order to ensure steady state expression of the reporter. We imaged cells on a Thermo Fisher CX7 LZR high content analysis platform set to 20x magnification,1×1 binning and applied illumination correction, resulting in a lateral pixel size of 207 nm. We collected z-stacks spanning 3 microns in 1 micron steps using 70 ms exposures of 488 nm light at 100% intensity (MCP-GFP), and 100 ms exposures of 561 nm light at 100% intensity (mCherry transfection reporter).

### Reporter transcription site image analysis

We projected image stacks in ImageJ (NIH) using Z Project (maximum intensity) ^59^, and split individual channels of each image for downstream processing. We created cell masks for each positively transfected cell from the mCherry signal using CellProfiler 4.0.7 to identify objects using global robust background thresholding ^60^. mCherry intensity was used to draw dividing lines between clumped objects. We discarded cell masks outside of a specified diameter range (50 - 200 px) or those touching the border of the image. We then cropped a region of interest around each individual cell mask using a custom MATLAB script to generate individual images each encompassing a single mCherry-positive cell. We detected individual transcription sites and measured their intensity using the spot detection software Airlocalize ^30^, resulting in spot counts and intensities per cell. For each condition, we defined the active fraction as the average number of detected TSs per cell, and the intensity as the average intensity of TSs across cells with detectable TSs. We collected data from two biological replicates for each condition and computed the error as the standard error of the mean.

### IDR feature and protein sequence analysis

For additional IDR and protein sequence features, we downloaded the canonical sequence of each protein from UniProt (or iGEM for VP64, http://parts.igem.org/Part:BBa_J176013). We extracted IDRs from protein sequences using metapredict with the metapredict-hybrid mode ^61^, with or without a threshold requiring that 60 or more residues of a protein are disordered (Table S3). This threshold ensures that sequence properties are calculated over a sufficient number of residues to be interpretable. Data presented in Figure S3A-B includes residues from all IDR data. Data presented in Figure S3C-D includes only IDR residues from proteins satisfying the 60+ threshold. When using IDR and protein sequence features as training data, full length protein sequences were used as inputs for feature calculation alongside the thresholded data with similar results. Data from full length protein sequences was ultimately used for model training as this dataset contained all CRISPRburst characterized TFs, compared to the reduced thresholded dataset.

We calculated metrics for the following features: length, amino acid composition, IDR sequence complexity, IDR fraction of charged residues, IDR net charge per residue, IDR kappa, IDR hydrophobicity (using Kyte-Doolittle hydrophobicity scale), IDR aromatic count, IDR aromatic + arginine count ^62^, number of low complexity sequences found in IDRs ^63^ (Table S3). Sequence properties were calculated using localCIDER ^64^. Sequences used for input and code used for analysis can be found on Github at https://github.com/holehouse-lab/supportingdata/tree/master/2022/mamrak_2022.

### Clustering

We clustered CRISPRburst-characterized TF data using unsupervised learning algorithms. Primarily, we applied Llyod’s algorithm (k-means clustering) on TF active fraction and intensity effects using 5 clusters, over 10 replicates. We determined the optimal cluster number to be 5 using the Elbow method, measuring the increase in variance explained (sum of intra-cluster distances per cluster) by an additional cluster (Figure S2A). We applied an orthogonal approach using principal component analysis (PCA) on all conditions in the dataset (Figure S2B). PCA identifies recruitment-specific active fraction and intensity effects (TF ABA / TF DMSO) as the major contributors to PC1 and PC2, respectively (Figure S2C), supporting the clustering analysis. Lastly, we clustered the data using density-based spatial clustering of application with noise (DBSCAN). DBSCAN parameters set to 0.15 for neighborhood search radius and 5 for minimal number of neighbors required for each core point identified both active fraction only TFs and intensity only TFs, consistent with the results of k-means clustering (Figure S2D).

### BioGRID data

We retrieved protein interaction data from the BioGRID database ^42^ version 4.4.207, and pruned it to include only physical interactions supported by experimental evidence. We filtered interaction pairs, keeping only those including a human transcription factor, based on a published TF database ^7^.

### Multivariate regression model

In order to select interactors to include in model training, we mined the list of pruned BioGRID interactions and selected proteins interacting with at least one of the 78 TF characterized by CRISPRburst, and at least one TF absent from the CRISPRburst dataset. Out of this set, we only included proteins expressed in HeLa cells, defined based on their RNA expression levels above a log(pTPM+1) threshold of 0.1 in a public RNA-seq dataset (Human Protein Atlas cell line data)^65^. Self-interactions were assigned to each TF equal to the maximum number of evidence that TF has for any factor. Interaction data was normalized per TFs and per interactors to avoid biases stemming from inherent differences between some TFs being well studied and having rich interaction data compared to those under-studied. We trained two partial least-squares regression models in MATLAB (plsregress) using matched interaction data and either the activity or intensity measured for characterized TFs. We further excluded interactors forming fewer than 240-258 interactions (for the active fraction model) or 30-45 interactions (for the intensity model). Four and five components were used for predictor variable principal component analysis, in the active fraction and intensity models respectively. The value of the interaction thresholds and the number of PCA components were determined by optimizing model performance on a test subset of TFs. The models output predict weights for each interactor protein; we applied those to interaction data for all non-characterized human TFs, resulting in separate active fraction and intensity predictions for each TF.

### ROC curves

We assigned each TF from the test set to its ground truth kinetic class based on whether its active fraction and intensity measured by CRISPRburst fell below or above threshold values (1.18 for intensity and 1.295 for active fraction). We chose these values because they are equidistant from cluster centroids obtained by k-means (Figure 2D), using 1) the low effect and intensity only clusters (for the intensity threshold) or 2) the low effect and active fraction only clusters (for the active fraction threshold). We then set arbitrary active fraction thresholds (resp. intensity) to classify the TF into a kinetic class based on its predicted active fraction (resp. predicted intensity). We gradually increased the values of the arbitrary thresholds and computed for each threshold a true positive rate (TPR) and false positive rate (FPR) by comparing the predicted kinetic class to the ground truth kinetic class.

### Flow cytometry analysis

We washed cells in DPBS and resuspended them in DPBS containing 1% paraformaldehyde. We measured GFP fluorescence with a Sony SH800S cell sorter (Sony Biotechnology, San Jose, CA). We applied software-based compensation during analysis using singly GFP positive and mCherry positive cells to determine spectral crosstalk between channels. We gated live cells according to forward and side scatter (singles), then drew a gate containing mCherry positive cells (mCherry+) using an untransfected control as reference. Within this gate, we defined GFP-positive cells (GFP+) using a fluorescence threshold determined by comparison with a no GFP control sample. We analyzed a minimum of 200,000 cells per condition. Figures presented throughout the manuscript are representative of two independent experiments.

## Acknowledgments

We thank NYU Langone’s Microscopy Laboratory (RRID: SCR_017934) staff for assistance with the Thermo Scientific CellInsight CX7 LZR high content analysis platform. This core is partially funded by NYU Cancer Center Support Grant NIH/NCI P30CA016087 and grant S10OD021727. This work was supported by NIH grants R01AG075272 (NEM & TL), R01GM127538 (NEM & TL), Melanoma Research Foundation Award 687306 (TL), NSF Graduate Research Fellowship DGE-1247271 (DG), CIHR Project Grant PJT-175277 (MT), and Longer Life Foundation grant (a collaboration between RGA and Washington University, ASH). We thank Edouard Bertrand (IGMM, Montpellier, France) for the gift of cell lines. This work uses cells derived from the HeLa cell line. Henrietta Lacks, and the HeLa cell line that was established from her tumor cells without her knowledge or consent in 1951, have made significant contributions to scientific progress and advances in human health. We are grateful to Henrietta Lacks, now deceased, and to her surviving family members for their contributions to biomedical research.

