## Supplementary Note and Figures for "The kinetic landscape of human transcription factors"

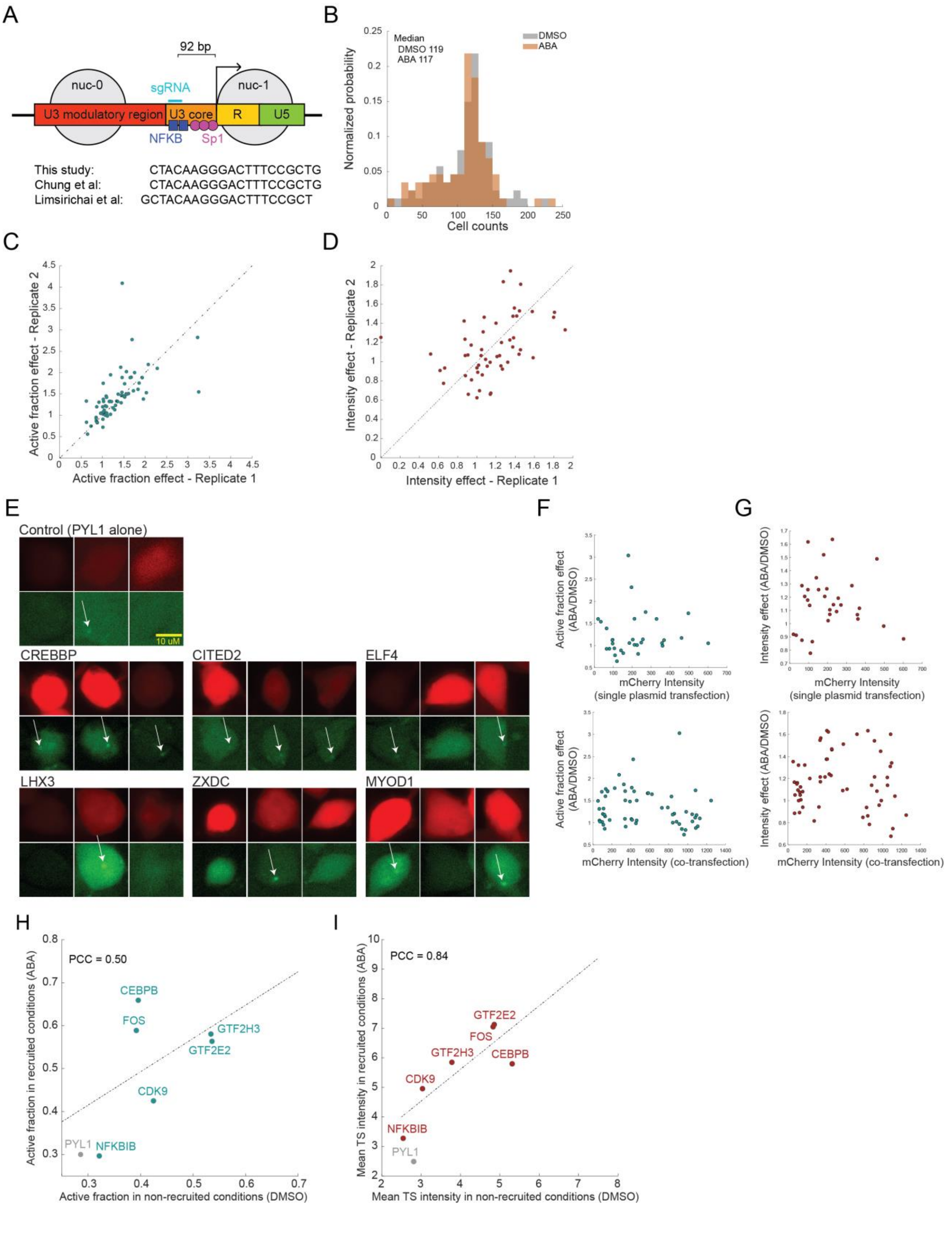

A
92 bp
nuc-0
nuc-1
sgRNA
U3 modulatory region
U3 core
R
U5
NFKB
Sp1
This study: CTACAAGGGACTTTCCGCTG
Chung et al: CTACAAGGGACTTTCCGCTG
Limsirichai et al: GCTACAAGGGACTTTCCGCT
B
Median
DMSO 119
ABA 117
DMSO
ABA
Normalized probability
Cell counts
C
Active fraction effect - Replicate 2
Active fraction effect - Replicate 1
D
Intensity effect - Replicate 2
Intensity effect - Replicate 1
E
Control (PYL1 alone)
10 uM
CREBBP
CITED2
ELF4
LHX3
ZXDC
MYOD1
F
Active fraction effect (ABA/DMSO)
mCherry Intensity (single plasmid transfection)
mCherry Intensity (co-transfection)
G
Intensity effect (ABA/DMSO)
mCherry Intensity (single plasmid transfection)
mCherry Intensity (co-transfection)
H
PCC = 0.50
CEBPB
FOS
GTF2H3
GTF2E2
CDK9
PYL1
NFKBIB
Active fraction in recruited conditions (ABA)
Active fraction in non-recruited conditions (DMSO)
I
PCC = 0.84
GTF2E2
FOS
GTF2H3
CEBPB
CDK9
NFKBIB
PYL1
Mean TS intensity in recruited conditions (ABA)
Mean TS intensity in non-recruited conditions (DMSO)

**Figure S1: Characterization of CRISPRburst framework**

**A** Location of the sgRNA binding site within the HIV-1 LTR. The sgRNA (teal) binds within a regulatory region (U3 core), and overlaps with a natural NF-κB binding site between nucleosome positions nuc-0 and nuc-1, 92 bp upstream of the transcription start site. sgRNA sequences from Chung *et al*. and Limsirichai *et al*. are shown for comparison [32, 33]. Black arrow designates the transcription start site. LTR schematic adapted from Limsirichai *et al*.

**B** Histograms showing the distribution of sample sizes across DMSO and ABA conditions. For each TF, we analyzed a median number of 119 (DMSO, gray), and 117 cells (ABA, orange) yielding 220 total cells per TF on average.

**C, D** Scatter plot comparing active fraction (**C**) and intensity (**D**) between biological replicates.

**E** Representative images of transfected cells upon TF recruitment. Top row shows the transfection marker, mCherry. The bottom row shows MCP-GFP. White arrows designate detected transcription sites. Scale bar: 10 μm.

**F**, **G** Scatter plots illustrating the relationship between fluorescent transfection marker, mCherry intensity and observed TF effect sizes for active fraction (**F**) and intensity (**G**). Experiments were performed using either one plasmid co-expressing TF-PYL1 and mCherry (top) or by co-transfecting separate plasmids, one expressing TF-PYL1 and one expressing mCherry (bottom).

**H, I** Scatter plot showing that 6 TFs which naturally bind the HIV LTR give rise to similar kinetics when expressed (DMSO) as they do upon dCas9 recruitment (ABA), demonstrating that synthetic recruitment reflects the kinetic trends observed in the natural context. **H:** active fraction; **I:** intensity. PYL1 shown for reference and was not included in correlation calculation.

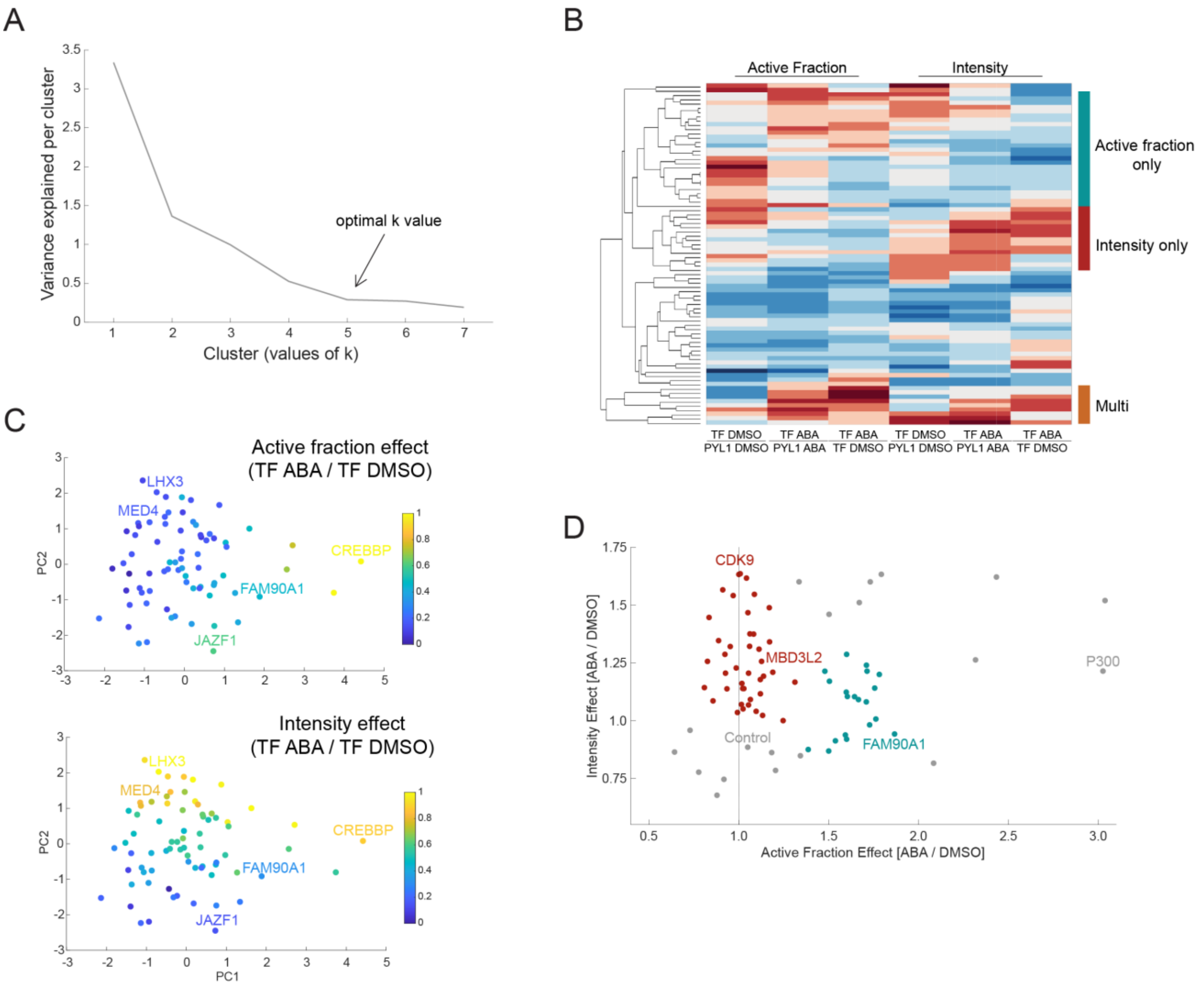
A
Variance explained per cluster
Cluster (values of k)
optimal k value
B
Active Fraction
Intensity
Active fraction only
Intensity only
Multi
TF DMSO PYL1 DMSO
TF ABA PYL1 ABA
TF ABA TF DMSO
TF DMSO PYL1 DMSO
TF ABA PYL1 ABA
TF ABA TF DMSO
C
Active fraction effect (TF ABA / TF DMSO)
LHX3
MED4
CREBBP
FAM90A1
JAZF1
PC2
Intensity effect (TF ABA / TF DMSO)
LHX3
MED4
CREBBP
FAM90A1
JAZF1
PC1
PC2
D
CDK9
MBD3L2
P300
Control
FAM90A1
Intensity Effect [ABA / DMSO]
Active Fraction Effect [ABA / DMSO]

**Figure S2: Additional analysis of characterized TF clusters**

**A** Variance explained per cluster in k-means analysis of CRISPRburst-characterized TFs. Variance explained, calculated as the sum of intra-cluster distances per cluster, decreases with increasing values of k. The optimal k value is determined as the point where increasing k no longer significantly increases the variance explained (k = 5).

**B** Heatmap showing the effects of TF expression alone (TF DMSO / PYL1 DMSO), recruitment alone (TF ABA / TF DMSO), or composite (TF ABA / PYL1 ABA), for active fraction and intensity measurements across all characterized TFs. Effects were clustered by TFs and presented as a column normalized clustergram.

**C** The 6 ratios per TF presented in **B** were subject to principal component analysis (PCA). Scatter plots of PC1 and PC2 are presented and colored active fraction effect (top), and intensity (bottom).

**D** Active fraction versus intensity for 78 characterized TFs colored by density-based spatial clustering (DBSCAN). Teal represents active fraction only TFs, red represents intensity only TFs, and gray represents other TFs.

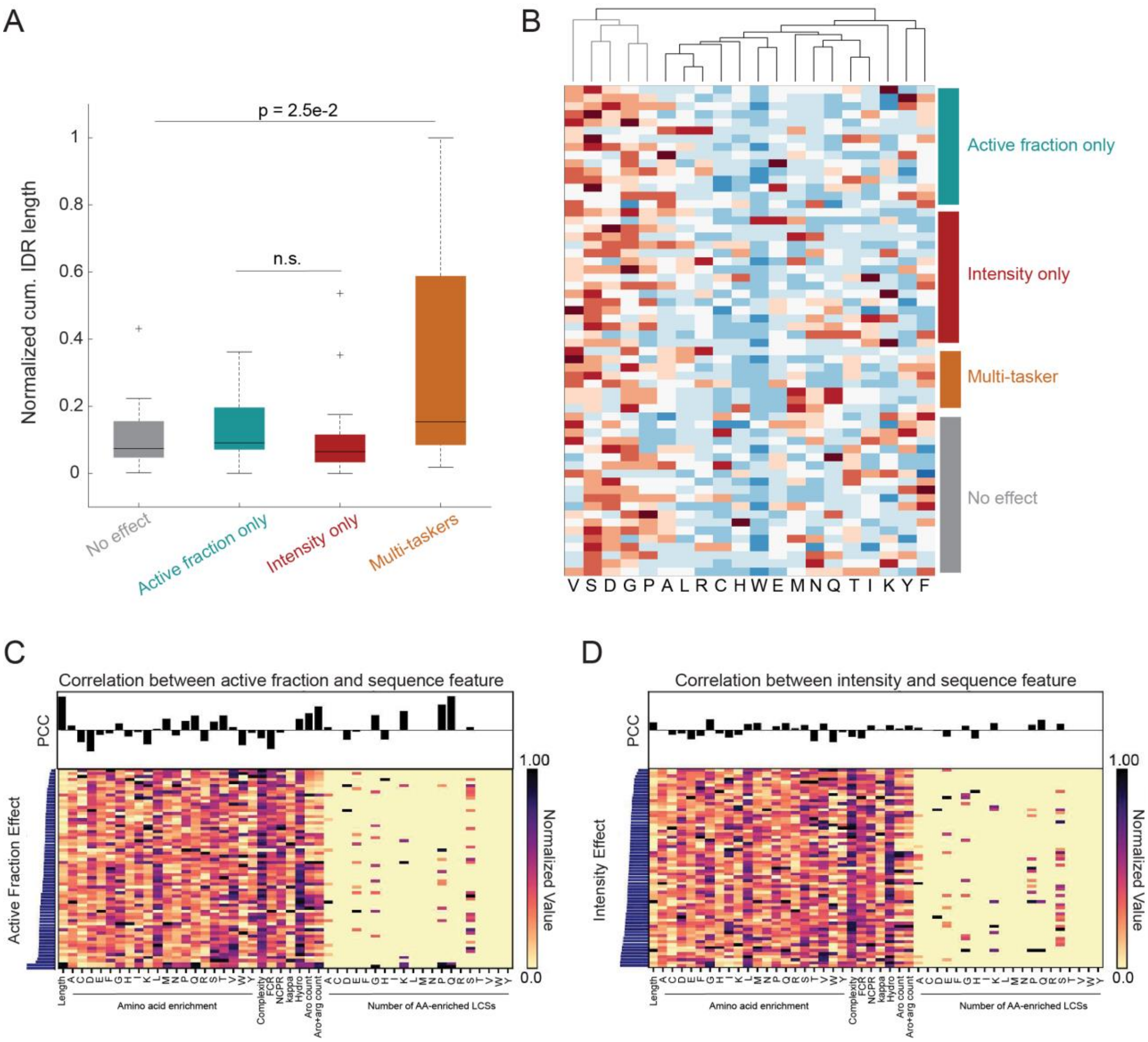

A
p = 2.5e-2
n.s.
Normalized cum. IDR length
1
0.8
0.6
0.4
0.2
0
No effect
Active fraction only
Intensity only
Multi-taskers
B
Active fraction only
Intensity only
Multi-tasker
No effect
V S D G P A L R C H W E M N Q T I K Y F
C
Correlation between active fraction and sequence feature
PCC
Active Fraction Effect
1.00
0.0
Normalized Value
Amino acid enrichment
Number of AA-enriched LCSs
D
Correlation between intensity and sequence feature
PCC
Intensity Effect
1.00
0.0
Normalized Value
Amino acid enrichment
Number of AA-enriched LCSs

**Figure S3: Intrinsically disordered region analysis of characterized TF clusters**

**A** Boxplot showing the cumulative length of IDRs per TF (normalized amino acid counts) for each cluster. On each box, the center mark indicates the median, and the bottom and top edges of the box indicate the 25th and 75th percentiles, respectively. Outliers are plotted individually as + symbols.

**B** Clustergram of amino acid frequencies per TFs in each cluster.

**C, D** Heatmaps illustrating the relationship between kinetic specialization and IDR features. Proteins were ranked in ascending order for either the active fraction (**C**) or intensity effect (**D**). Normalized matrices of 48 features (See methods) are presented as heat maps, where the color reflects the rank-ordering of smallest value (yellow) to large value (black). PCC between active fraction or intensity effect with each feature is provided above.

Significance was determined by two sample t-tests.

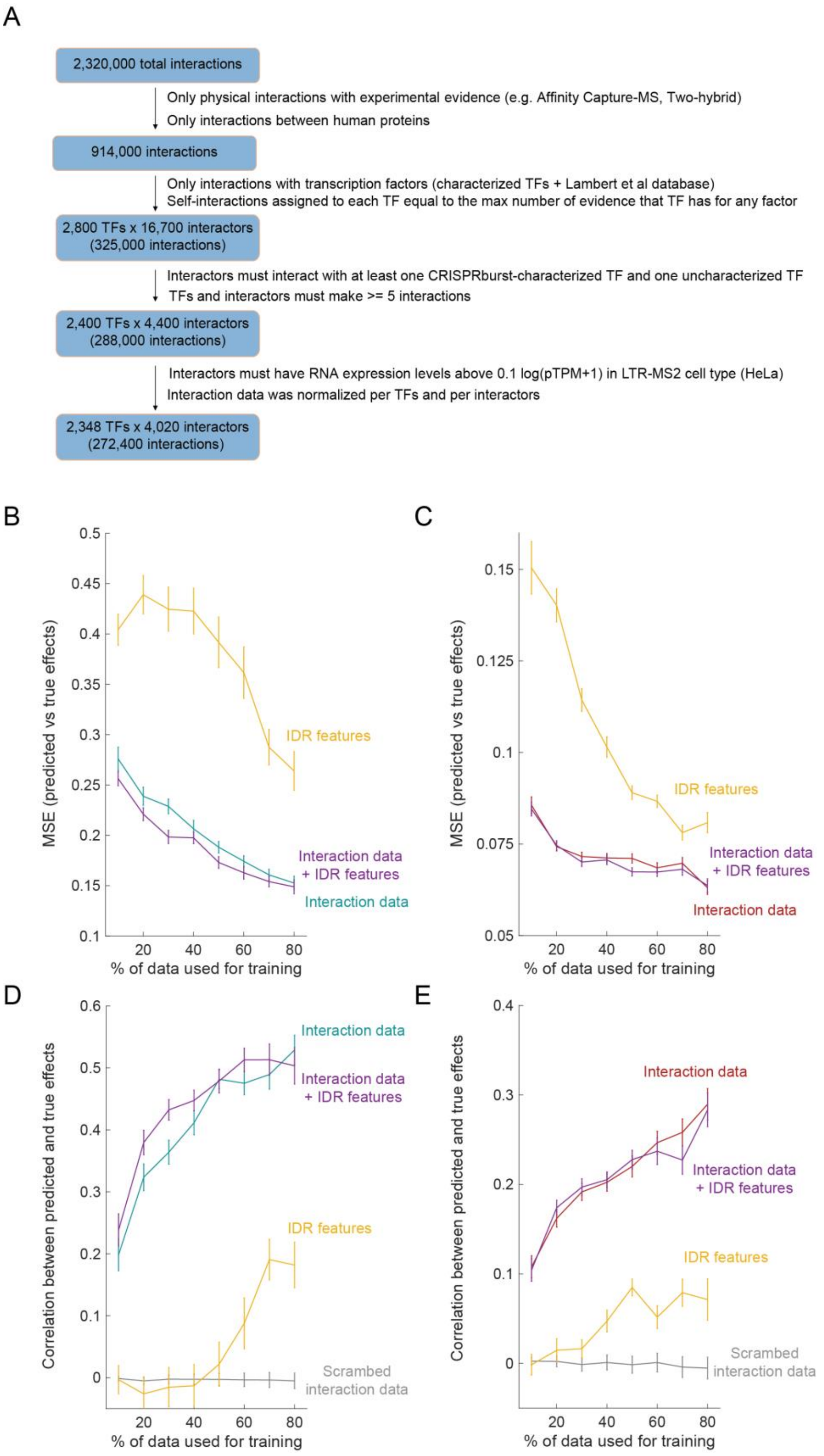
A
2,320,000 total interactions
Only physical interactions with experimental evidence (e.g. Affinity Capture-MS, Two-hybrid)
Only interactions between human proteins
914,000 interactions
Only interactions with transcription factors (characterized TFs + Lambert et al database)
Self-interactions assigned to each TF equal to the max number of evidence that TF has for any factor
2,800 TFs x 16,700 interactors
(325,000 interactions)
Interactors must interact with at least one CRISPRburst-characterized TF and one uncharacterized TF
TFs and interactors must make >= 5 interactions
2,400 TFs x 4,400 interactors
(288,000 interactions)
Interactors must have RNA expression levels above 0.1 log(pTPM+1) in LTR-MS2 cell type (HeLa)
Interaction data was normalized per TFs and per interactors
2,348 TFs x 4,020 interactors
(272,400 interactions)
B
MSE (predicted vs true effects)
% of data used for training
IDR features
Interaction data
+ IDR features
Interaction data
C
MSE (predicted vs true effects)
% of data used for training
IDR features
Interaction data
+ IDR features
Interaction data
D
Correlation between predicted and true effects
% of data used for training
Interaction data
Interaction data
+ IDR features
IDR features
Scrambed
interaction data
E
Correlation between predicted and true effects
% of data used for training
Interaction data
Interaction data
+ IDR features
IDR features
Scrambed
interaction data

**Figure S4: Characterization of the multivariate regression model**

**A** Flow chart describing the processing of BioGRID data from the full interaction database down to the interaction matrix used for training and prediction of TF function.

**B, C** Mean squared error (MSE) between model predictions and measured kinetics for active fraction (**B**) and intensity (**C**), as a function of increasing training set size. Curves represent results from a model trained on BioGRID interaction data (teal or red), IDR features (yellow), interaction data and IDR features combined (purple). Each value is the average of 100 MSE calculations computed from randomly assigned training/validation sets.

**D, E** Mean Pearson’s correlation coefficient between model predictions and measured kinetics for active fraction (**D**) and intensity (**E**), as a function of increasing training set size. Curves represent results from a model trained on the real interaction matrix (teal or red), a randomly scrambled interaction matrix (gray), IDR features (yellow), or interaction data and IDR features combined (purple). Each value is the average of 100 PCCs computed from randomly assigned training/validation sets.

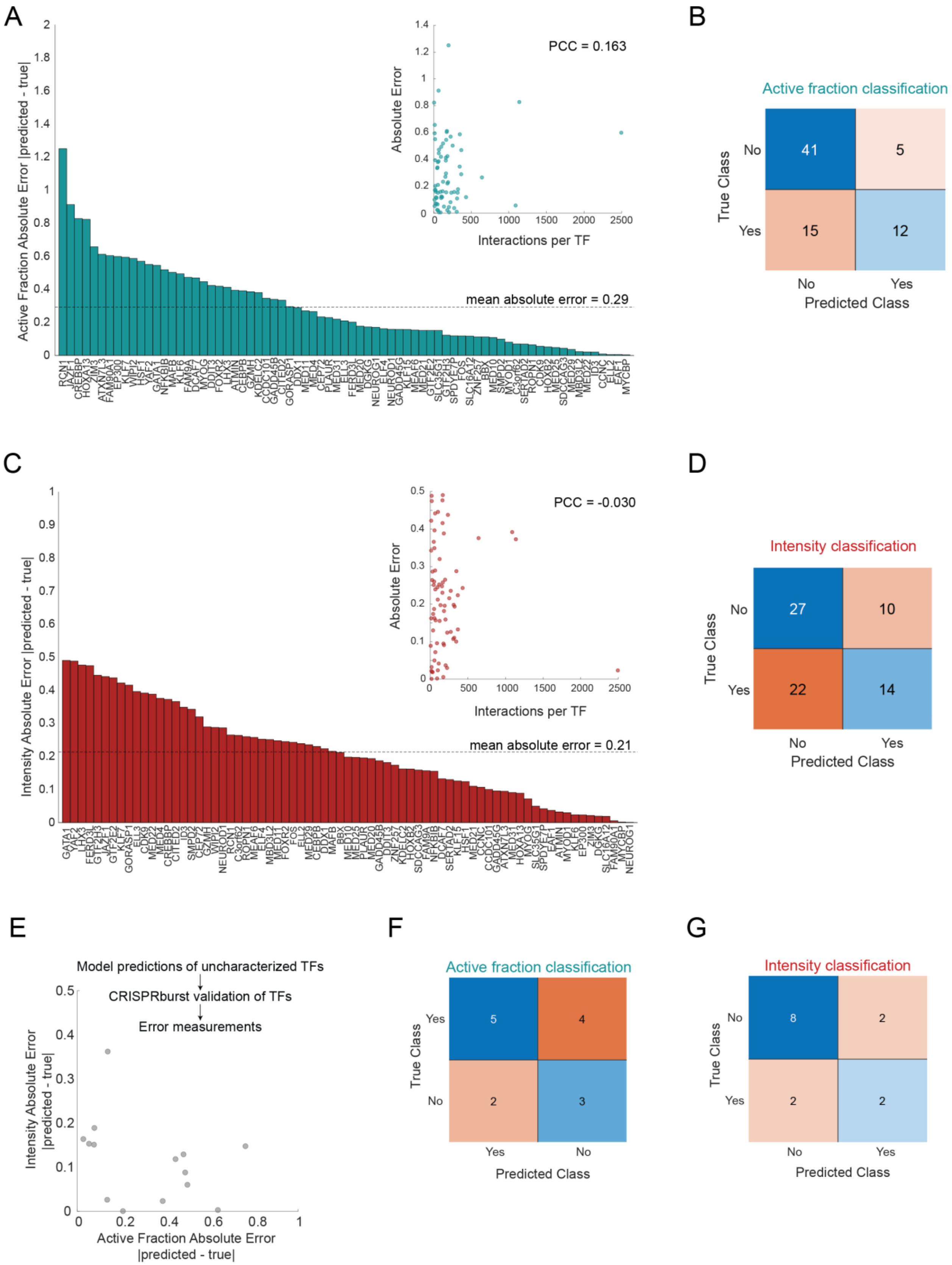
A
Active Fraction Absolute Error |predicted - true|
mean absolute error = 0.29
PCC = 0.163
Absolute Error
Interactions per TF
B
Active fraction classification
True Class
No
Yes
41
5
15
12
No
Yes
Predicted Class
C
Intensity Absolute Error |predicted - true|
mean absolute error = 0.21
PCC = -0.030
Absolute Error
Interactions per TF
D
Intensity classification
True Class
No
Yes
27
10
22
14
No
Yes
Predicted Class
E
Model predictions of uncharacterized TFs
CRISPRburst validation of TFs
Error measurements
Intensity Absolute Error |predicted - true|
Active Fraction Absolute Error |predicted - true|
F
Active fraction classification
True Class
Yes
No
5
4
2
3
Yes
No
Predicted Class
G
Intensity classification
True Class
No
Yes
8
2
2
2
No
Yes
Predicted Class

**Figure S5: Performance of the multivariate regression model for the prediction of TF kinetic classes.**

**A, C** Ranked bar graph depicting the absolute error in active fraction (**A)** or intensity (**C**) per characterized TF when training on the full dataset except for the TF being predicted. The black dashed line represents the mean error across the dataset. Insets show the absolute error in active fraction (**A**) or intensity (**C**) as a function of interaction count per characterized TF.

**B, D** Confusion matrix presenting the incidence of true positives, false positives, false negatives, and true negatives when classifying factors into active fraction (**B**) and intensity (**D**) classes based on the predicted effects of cross validated training data compared to the ground truth effects measured by CRISPRburst, and applying thresholds based on k-means clustering.

**E** Absolute error between bursting kinetics predicted by the model and values measured by CRISPRburst for a test set of 14 TFs not previously seen by the model. Mean squared errors: 0.15 (active fraction), 0.025 (intensity).

**F, G** Confusion matrix presenting the incidence of true positives, false positives, false negatives, and true negatives when classifying factors into active fraction (**F**) intensity (**G**) classes based on the predicted effects of  14 TFs never seen by the model compared to the ground truth effects measured by CRISPRburst, and applying thresholds based on k-means clustering.

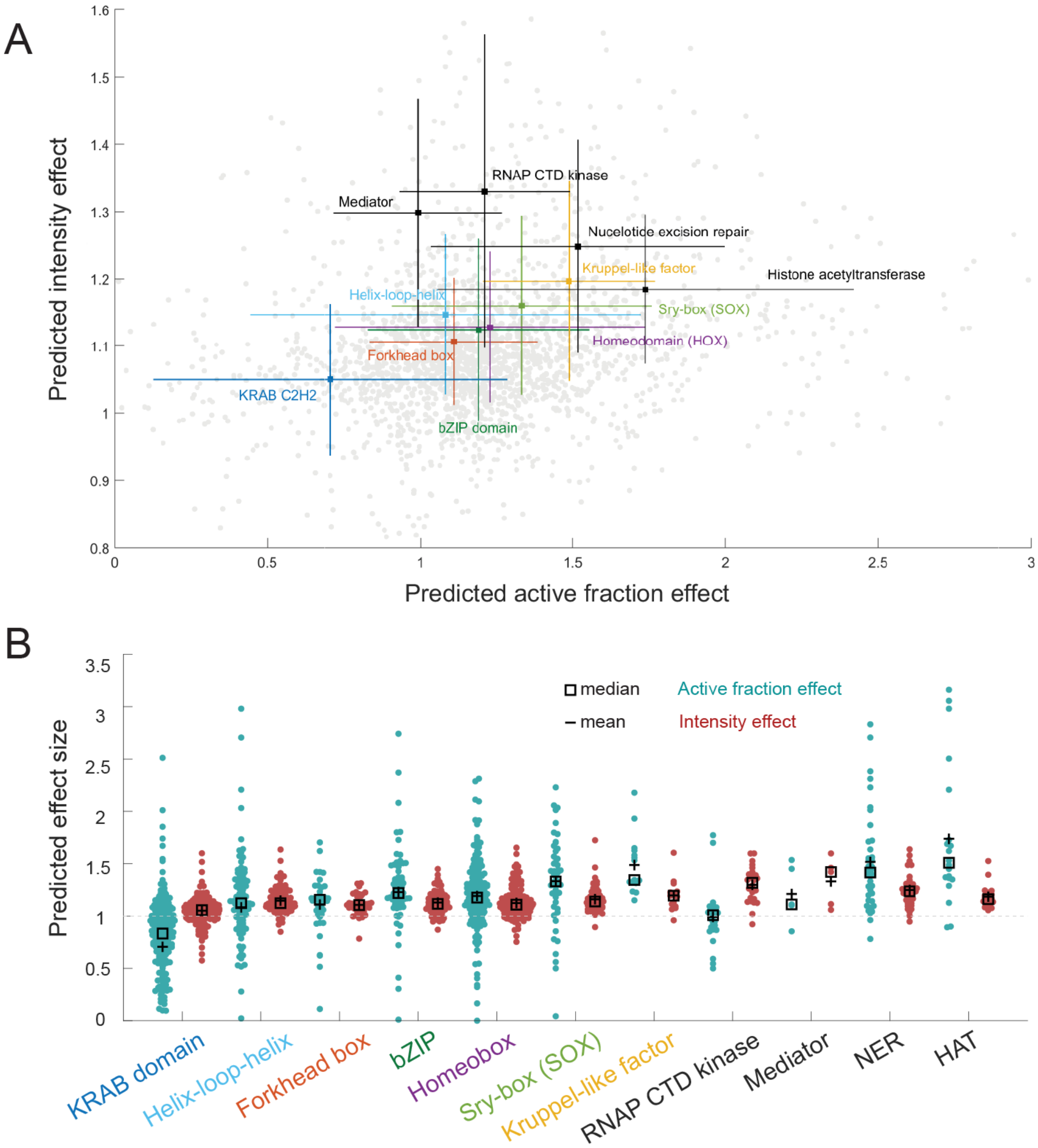
A
Predicted intensity effect
Predicted active fraction effect
0
0.5
1
1.5
2
2.5
3
0.8
0.9
1
1.1
1.2
1.3
1.4
1.5
1.6
RNAP CTD kinase
Mediator
Nucelotide excision repair
Kruppel-like factor
Histone acetyltransferase
Helix-loop-helix
Sry-box (SOX)
Homeodomain (HOX)
Forkhead box
KRAB C2H2
bZIP domain
B
Predicted effect size
0
0.5
1
1.5
2
2.5
3
3.5
median
mean
Active fraction effect
Intensity effect
KRAB domain
Helix-loop-helix
Forkhead box
bZIP
Homeobox
Sry-box (SOX)
Kruppel-like factor
RNAP CTD kinase
Mediator
NER
HAT

**Figure S6: TF kinetics are widely variable within TF families.**

**A** Scatter plot illustrating the variability between classically defined TF families[7]. Predicted kinetic signatures for individual TF families are plotted as crosses centered on the average kinetic metric per family; the length of the cross arms represents the standard deviation within each family. Co-activating complexes (gray) are plotted for reference. RNAP CTD: RNA Polymerase II carboxy-terminal domain.

**B** Same as A, represented as swarm plots for each TF family. Active fraction effects are plotted in teal, intensity effects are plotted in red. The mean and median effects per family are represented by black dashes and black squares, respectively.

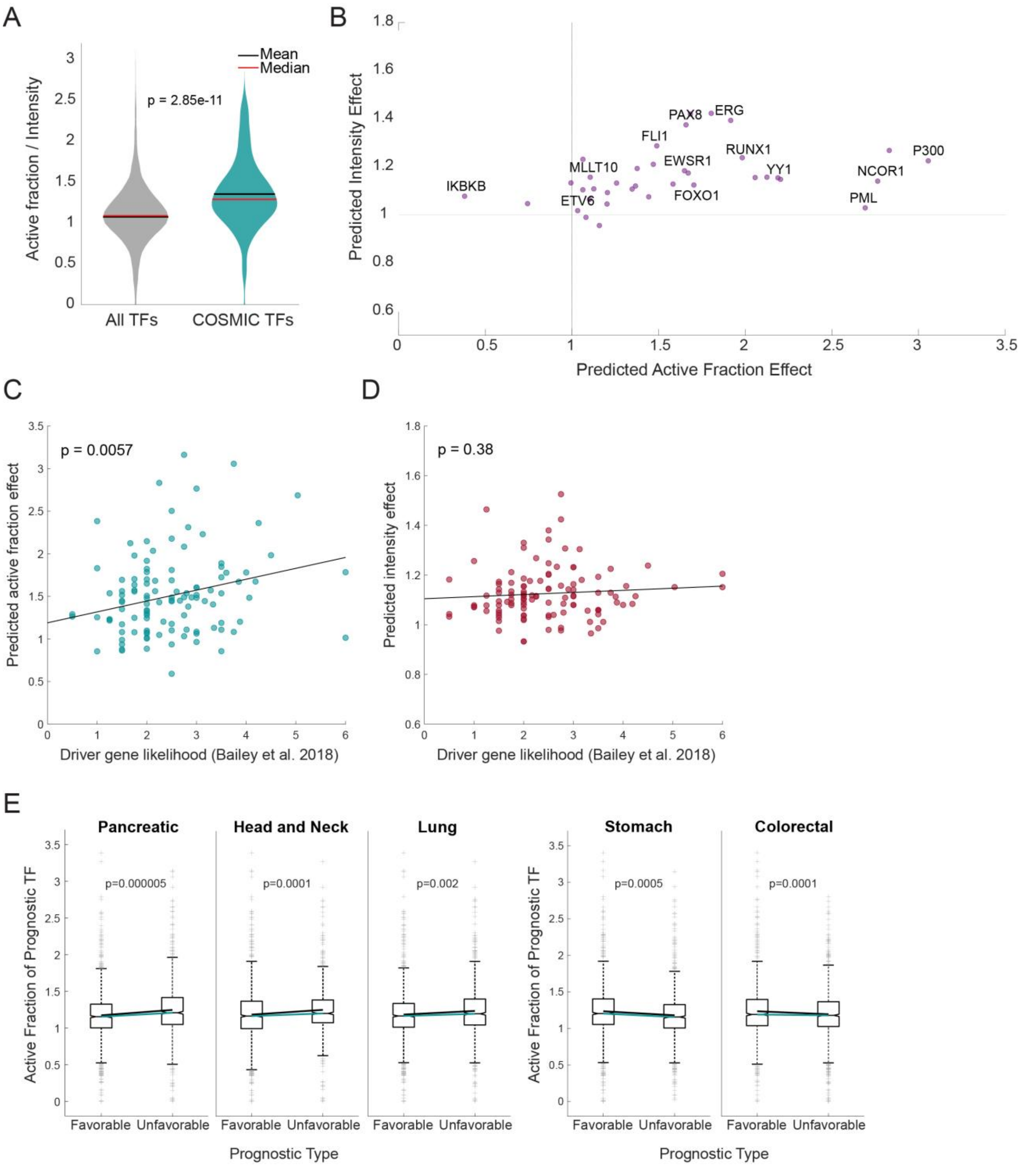

A
Mean
Median
p = 2.85e-11
Active fraction / Intensity
All TFs
COSMIC TFs
B
Predicted Intensity Effect
Predicted Active Fraction Effect
IKBKB
MLLT10
ETV6
FLI1
PAX8
ERG
EWSR1
FOXO1
RUNX1
YY1
P300
NCOR1
PML
C
p = 0.0057
Predicted active fraction effect
Driver gene likelihood (Bailey et al. 2018)
D
p = 0.38
Predicted intensity effect
Driver gene likelihood (Bailey et al. 2018)
E
Pancreatic
Head and Neck
Lung
Stomach
Colorectal
p=0.000005
p=0.0001
p=0.002
p=0.0005
p=0.0001
Active Fraction of Prognostic TF
Favorable
Unfavorable
Prognostic Type

**Figure S7: Cancer drivers are enriched for TFs increasing their target active fraction.**

**A** Violin plots of the ratio of predicted active fraction relative to intensity for all TFs (gray) compared to TFs within the catalog of somatic cancer mutations (COSMIC, teal) [48]. Significance was determined by two sample t-tests.

**B** TFs commonly found in oncogenic transcription factor fusion genes (purple) are enriched in high active fraction kinetics relative to all human TFs (gray) [49].

**C, D** Scatter plot illustrating the relationship between driver gene likelihood (defined in Bailey *et al*. 2018) and the predicted active fraction (**C**) or predicted intensity (**D**) for TFs within the COSMIC database [48,50].

**E** Boxplots showing the predicted active fraction effect of TFs which high expression carries a favorable vs. poor prognosis in various cancer types. Prognostic expression data from Uhlen *et al.* 2017[51]. On each box, the center mark indicates the median, and the bottom and top edges of the box indicate the 25th and 75th percentiles, respectively. Outliers are plotted individually as + symbols. 5/17 cancer types exhibit statistical differences for the predicted active fraction between prognosis TF groups. We did not observe any statistically significant difference between prognosis groups for the predicted intensity.

Significance was determined by two sample t-tests, correcting for multiple hypothesis testing.

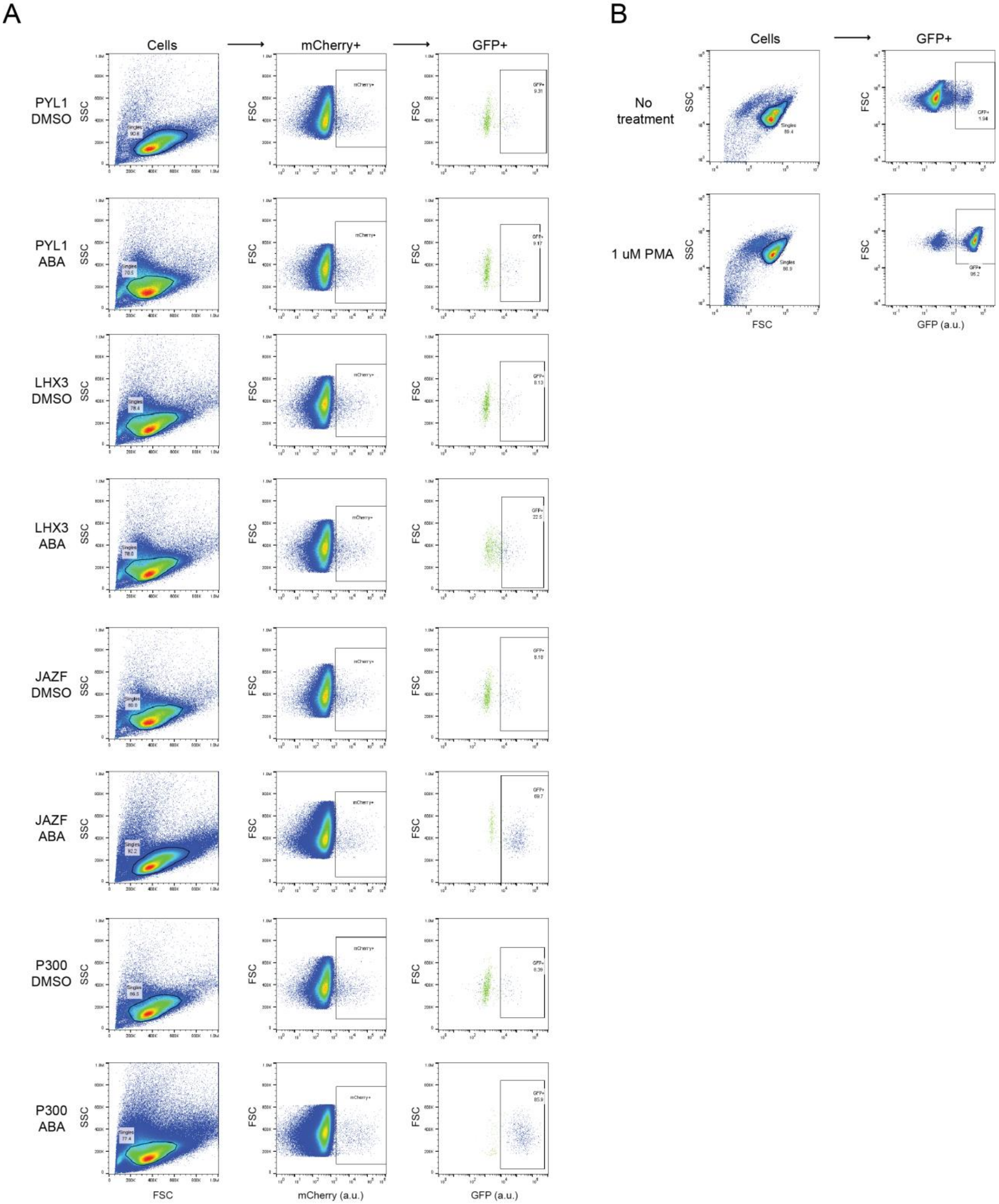

A
Cells
mCherry+
GFP+
PYL1 DMSO
PYL1 ABA
LHX3 DMSO
LHX3 ABA
JAZF DMSO
JAZF ABA
P300 DMSO
P300 ABA
SSC
FSC
mCherry (a.u.)
GFP (a.u.)
B
Cells
GFP+
No treatment
1 uM PMA
SSC
FSC
GFP (a.u.)

**Figure S8: FACS gates used in J-Lat activation experiments.**

**A** FACS plots demonstrating the gating strategy for quantifying GFP+ cell frequencies. Cells were gated on forward scatter (FSC) and side scatter (SSC) to separate single cells (left column). We then isolated transfected cells based on expression of the mCherry transfection marker (central column). We set mCherry gate using an un-transfected sample as reference. Within the gated mCherry+ population, we eventually quantified the fraction of GFP+ cells. We set the GFP+ gate using both a negative (untreated J-Lat cells) and positive control (PMA treated J-Lat cells) to establish the upper and lower limits of the GFP signal. A minimum of 215,000 cells were analyzed per sample.

**B** FACS plots for negative (untreated J-Lat cells) and positive control (J-Lat cells treated with 1 μM PMA). 100,000 cells were analyzed per sample.

**Table S1: TF characterization data generated using CRISPRburst**

**Table S2: Gene ontology enrichments for characterized TFs by cluster**

**Table S3: IDR and protein sequence features for characterized TFs**

**Table S4: Model validation using CRISPRburst on 14 uncharacterized TFs**

**Table S5: Predicted kinetic effects for all human TFs**

**Table S6: Gene ontology enrichments for most predictive interactors**

**Note 1: Image analysis pipeline for CRISPRburst data**

**Note 2: Description of interaction data processing and model construction**

### Note 1

#### Selecting for mCherry positive cells using CellProfiler 4.0.7

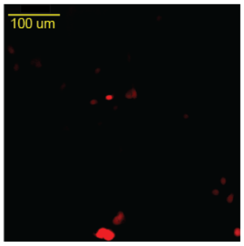


- Create list of files to be analyzed
- Identify Primary Objects
  - Supply typical size range for objects of interest
    - 50 – 200 pixels, for example
  - Pick thresholding method
    - Robustbackground
- Objects to Image
  - File type: uint16
- Save Images

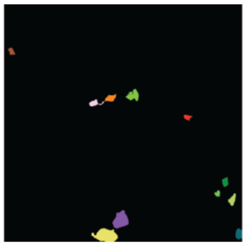

#### Selecting and cropping single cells from masked image in Matlab

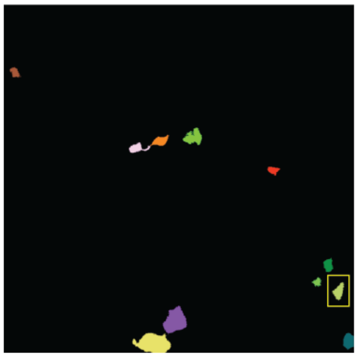

- Inputs: Folder full of mask images, each with a collection of objects
  - Each cell with its own ID (i.e. intensity value)
- For each input image, the script loops through objects, crops them, and saves in own image per each.
- Save mCherry and GFP images

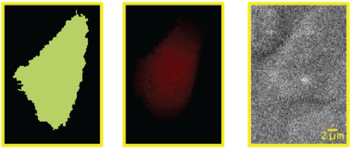


#### Spot detection from cropped cells using airlocalize in Matlab

- Airlocalize using batch mode per well
  - "contains A02" for example
- Can complete entire column if the same conditions are imaged within that column.
- Settings for AIRLOCALIZE(); are used for single time point analysis
  - TS intensity and active fraction effect calculations from position and intensity outputs per spot

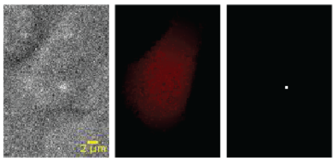


**sigma_xy: 2.0786** → Spot shape
sigma_z: 2
dx: 64
dy: 64
dz: 200
cutsize: 3
tol: 0.01
thickness: 1
**thresh.level: 6**
**thresh.units: SD** } → Spot intensity
thresh.sd: 11.2796

**Note 2**

Four datasets are required for TF kinetic predictions:

1. BioGRID database .tab3 formatted file version 4.4.207, downloaded from BioGRID file repository (https://downloads.thebiogrid.org/BioGRID)
2. Bursting data generated by CRISPRburst imaging, burstingData.txt file available on GitHub.
3. Full human TF database from Lambert *et al* (PMID:29425488, Document S1, Table S1), downloaded as .txt formatted file from http://humantfs.ccbr.utoronto.ca/download.php
4. RNA-seq data for HeLa cell line (RNA HPA cell line gene data, rna_celline.tsv, filtered on “Cell line” column to include only “HeLa” entries), from a public RNA-seq dataset via Human Protein Atlas cell line data (PMID:33149304).

BioGRID database was downloaded from the BioGRID file repository (https://downloads.thebiogrid.org/BioGRID) as a .tab3 formatted file.

The analysis pipeline consists of the following steps. The names in blue correspond to separate functions of the script, available on GitHub, https://github.com/timotheelionnet/burstingPrediction.

**Data Import**

The BioGRID data is used to generate a list of all interactions between all transcription factors: from the .tab3 format BioGRID downloaded file, all physical interactions between human proteins are collected and output as a 2 column list, where the first column is the gene symbol of the bait and the second column is the gene symbol of the hit. The same bait/hit gene pair might appear multiple times in the list if there are multiple evidences for that physical interaction in BioGRID (i.e. the bait/hit gene pair appears in distinct datasets curated within BioGRID).

**Building and pruning the Interaction Matrix**

The 2-column interaction list is pruned to include only interactions where at least one partner is a transcription factor. Proteins are called Transcription Factors if they are as either members of the Lambert *et al* TF database, and/or have been characterized by CRISPRburst.

An interaction matrix with n rows (corresponding to the number of TFs) and m columns (corresponding to the number of interactors) is initialized with zeros, and then populated with the number of evidences for each TF-interactor pair present in the pruned list. Each TF is set to interact with itself a number of times equal to the maximum number of evidence that the TF has for any other interactor.

The interaction matrix is pruned to remove interactors that do not make interactions with at least one CRISPRburst-characterized TF and at least one non-characterized TF (this is necessary to provide training information relevant to predictions outside the characterized set of TFs). TFs and interactors that make fewer than 5 interactions in total are also removed from the matrix.

**Selection of genes based on expression in HeLa cells**

Interactor genes are classified based on their expression in HeLa cells. Using RNA-seq data for HeLa cells (from a public RNA-seq dataset via Human Protein Atlas cell line data), all genes with RNA expression levels above a log(pTPM+1) threshold of 0.1 are considered expressed in HeLa cells. Interactors under the threshold are excluded from the model and the corresponding columns deleted from the interaction matrix.

At this step the interaction matrix is finished being pruned for training. The following plots describing the pruned interaction dataset:

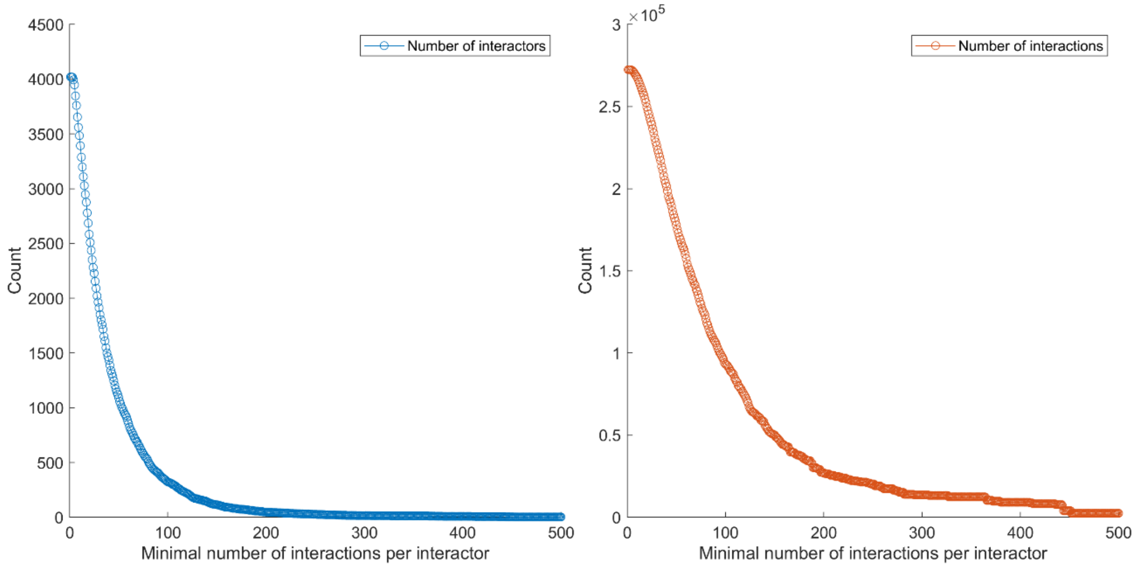


**Left:** Number of Interactors forming at least n_interactions total interactions as a function of n_interactions. **Right:** Total number of interactions in the matrix when keeping only interactors forming at least n_interactions total interactions , as a function of n_interactions; During pruning, only the TFs forming at least 5 interactions are retained in the interaction matrix.

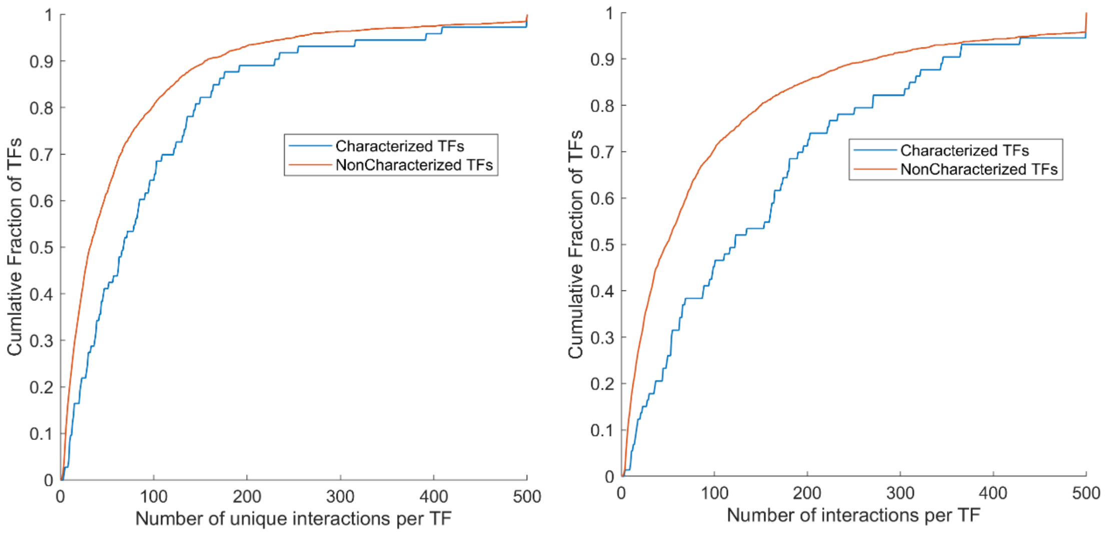

**Left:** Cumulative probability distribution of the number of unique interactions made by TFs characterized by CRISPRburst (blue) or non-characterized TFs (red); **Right:** Cumulative probability distribution of the number of total interactions made by TFs characterized by CRISPRburst (blue) or non-characterized TFs (red).

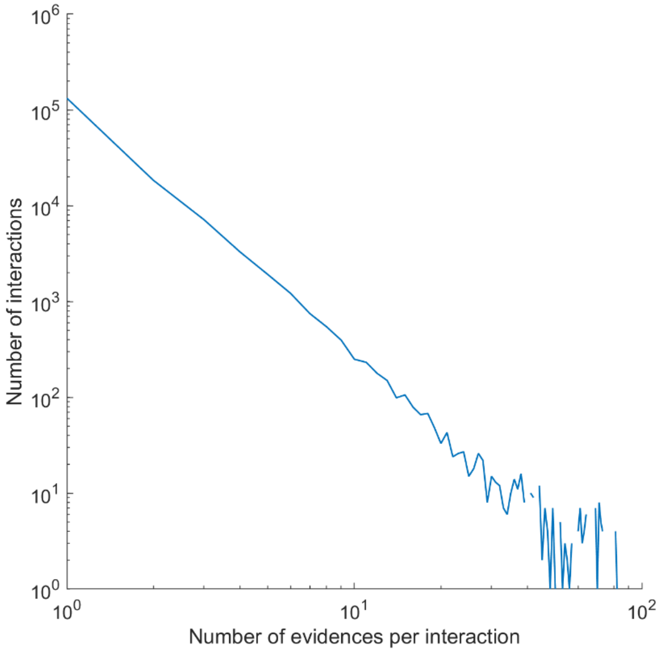


Distribution of the frequency of evidences per interaction pair.

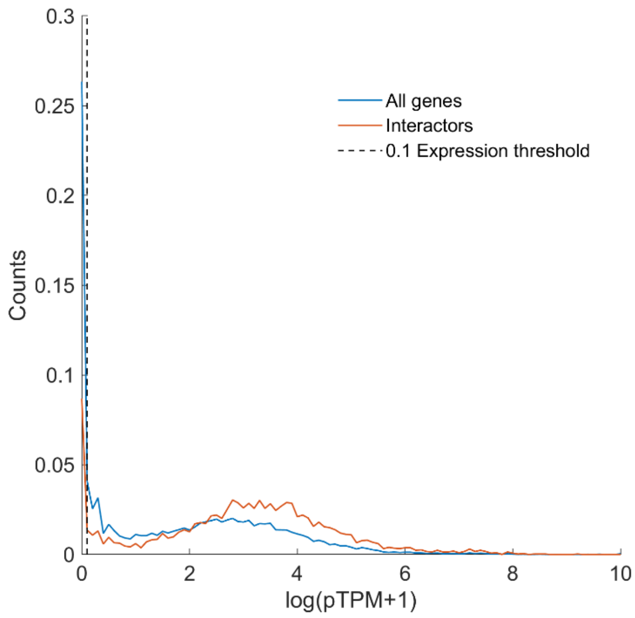


Distribution of RNA expression levels in HeLa cells for all genes (blue) and for genes selected as interactors before gating for expression (red). A dashed line at 0.1 log(pTPM+1) denotes the minimum expression level required to be included in the interaction matrix.

**Normalization of the Interaction Matrix**

In order to avoid biases between highly studied TFs (exhibiting exhaustive interaction data in BioGRID) and under-studied TFs (with sparse BioGRID interactions), we normalize the interaction matrix. First the interactions of each TF are normalized (z-score transform across each row); second, those TF-normalized interactions are z-score transformed per interactor (z-score transform across each column). We verified that other normalization strategies (e.g. dividing each TF interactions by the maximum evidence observed across interactors, or the sum of evidences observed for all interactors) did not substantially impact the model results.

**Building the regression model**

We then train the regression model on a training set drawn from TFs characterized by CRISPRburst.
First, we prune interactors separately for the models predicting active fraction and intensity, keeping only in the training the interactors making more than a minimum number of interactions overall.
Then we train two Partial Least-Squares (PLS) regression models using either the activity or intensity measured for characterized TFs and the matched TF interaction data.
Additional regression approaches were explored including support vector machine (SVM) regression and generalized additive model (GAM) regression. Tree based random forest classifiers were also tested for assigning TFs into clusters based on their predicted kinetic signatures. These approaches yielded similar performance metrics and identified similar interactors as most predictive, however due to the direct interpretability of linear regression models while achieving good performance metrics, we chose PLS regression.
The hyperparameters (interaction thresholds and number of PLS components) used for each model are obtained from optimization across a range of values, using cross validation with characterized TF data. We use four and five components for PLS component for active fraction and intensity modeling respectively; minimum number of interactions per interactor of 253 and 40 for active fraction and intensity modeling respectively.

**Predictions of TF kinetic signatures**

We apply the weights assigned by the models to each interactor protein to predict kinetic signatures for each non-characterized TF based on its interaction data. This results in a predicted effect for both active fraction effect and intensity effect separately. We also extract the 100 top predictors for each kinetic metric using beta values calculated by models for active fraction and intensity fraction.

In order to assess the robustness of the model to hyperparameter changes, we calculate predicted effects across a range of interaction thresholds (-5 to +5, centered on the optimal thresholds of 253 and 40 for active fraction and intensity). We then compute the standard deviation of each predicted kinetic metric as a measurement of prediction stability across an interaction range.

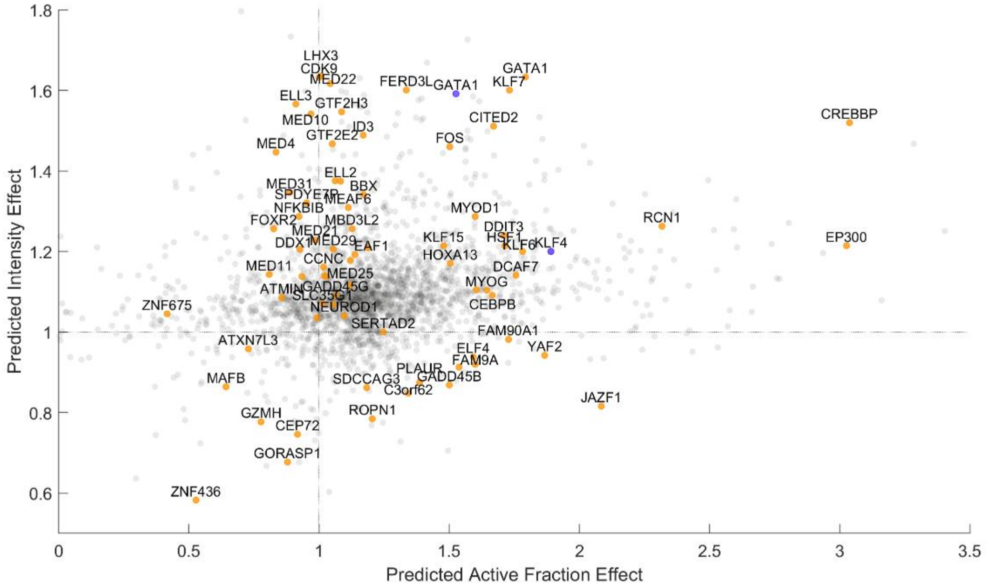


Model output: Predicted active fraction effect and intensity effect for all TFs. Each predicted TF is plotted in gray with the characterized TFs from CRISPRburst used as training data plotted in orange. Predictions for arbitrary examples overlaid in blue.
